# Common DNA sequence variation influences 3-dimensional conformation of the human genome

**DOI:** 10.1101/592741

**Authors:** David U. Gorkin, Yunjiang Qiu, Ming Hu, Kipper Fletez-Brant, Tristin Liu, Anthony D. Schmitt, Amina Noor, Joshua Chiou, Kyle J Gaulton, Jonathan Sebat, Yun Li, Kasper D. Hansen, Bing Ren

## Abstract

The 3-dimensional (3D) conformation of chromatin inside the nucleus is integral to a variety of nuclear processes including transcriptional regulation, DNA replication, and DNA damage repair. Aberrations in 3D chromatin conformation have been implicated in developmental abnormalities and cancer. Despite the importance of 3D chromatin conformation to cellular function and human health, little is known about how 3D chromatin conformation varies in the human population, or whether DNA sequence variation between individuals influences 3D chromatin conformation. To address these questions, we performed Hi-C on Lymphoblastoid Cell Lines (LCLs) from 20 individuals. We identified thousands of regions across the genome where 3D chromatin conformation varies between individuals and found that this conformational variation is often accompanied by variation in gene expression, histone modifications, and transcription factor (TF) binding. Moreover, we found that DNA sequence variation influences several features of 3D chromatin conformation including loop strength, contact insulation, contact directionality and density of local *cis* contacts. We mapped hundreds of Quantitative Trait Loci (QTLs) associated with 3D chromatin features and found evidence that some of these same variants are associated at modest levels with other molecular phenotypes as well as complex disease risk. Our results demonstrate that common DNA sequence variants can influence 3D chromatin conformation, pointing to a more pervasive role for 3D chromatin conformation in human phenotypic variation than previously recognized.

## INTRODUCTION

3-dimensional (3D) organization of chromatin is essential for proper regulation of gene expression^1–3^, and plays an important role in other nuclear processes including DNA replication^4,5^, X chromosome inactivation^6–9^, and DNA repair^10,11^. Many recent insights about 3D chromatin conformation have been enabled by a suite of technologies based on Chromatin Conformation Capture (3C)^12^. A high-throughput version of 3C called “Hi-C” enables the mapping of 3D chromatin conformation at genome-wide scale^13^, and has revealed several key features of 3D chromatin conformation including: 1) compartments (often referred to as “A/B compartments”), which refer to the tendency of loci with similar transcriptional activity to physically segregate in 3D space^13–15^, 2) chromatin domains (often referred to as Topologically Associating Domains, or TADs) demarcated by sharp boundaries across which contacts are relatively infrequent^16–18^, 3) chromatin loops, which describe point-to-point interactions that occur more frequently than would be expected based on the linear distance between interacting loci, and often anchored by convergent CTCF motif pairs^14^, and 4) Frequently Interacting Regions (FIREs), which are regions of increased local interaction frequency enriched for tissue-specific genes and enhancers^19,20^.

Previous studies have used Hi-C to profile 3D chromatin conformation across different cell types^14,16,21^, different primary tissues^19^, different cell states^22^, and in response to different genetic and molecular perturbations^23–27^, producing a wealth of knowledge about key features of 3D chromatin conformation. However, to our knowledge no study to date has measured variation in 3D chromatin conformation across more than a handful of unrelated individuals. Several observations demonstrate that at least in some cases DNA sequence variation between individuals can alter 3D chromatin organization with pathological consequences^28^. Pioneering work by Mundlos and colleagues described several cases in which rearrangements of TAD structure lead to gene dysregulation and consequent developmental malformations^29,30^. In cancer, somatic mutations and aberrant DNA methylation can disrupt TAD boundaries leading to dysregulation of proto-oncogenes^31,32^. Moreover, many genetic variants associated with human traits by GWAS occur in distal regulatory elements that loop to putative target gene promoters in 3D, and in some cases, the strength of these looping interactions has been shown to vary between alleles of the associated SNP^33,34^. Although these studies demonstrate that that both large effects as well as more subtle aberrations of 3D chromatin conformation are potential mechanisms of disease, population-level variation in 3D chromatin conformation more broadly has remained unexplored.

In the present study, we set out to characterize inter-individual variation in 3D chromatin conformation by performing Hi-C on Lymphoblastoid Cell Lines (LCLs) derived from individuals whose genetic variation has been cataloged by the HapMap or 1000 Genomes Consortia^35^. LCLs have been used as a model system to study variation in several other molecular phenotypes including gene expression, histone modifications, transcription factor (TF) binding, and chromatin accessibility^36–42^. These previous efforts provide a rich context to explore variation in 3D chromatin conformation identified in this model system. Through integrative analyses, we found that inter-individual variation in 3D chromatin conformation occurs on many levels including compartments, TAD boundary strengths, FIREs, and looping interaction strengths. Moreover, we found that variation in 3D chromatin conformation coincides with variation in activity of the underlying genome sequence as evidenced by transcription, histone modifications, and TF binding. Although our sample size is small, we observe reproducible effects of DNA sequence variation on 3D chromatin conformation and identify hundreds of Quantitative Trait Loci (QTLs) associated with multiple features of 3D chromatin conformation. Our results demonstrate that variation in 3D chromatin conformation is readily detectable from Hi-C data, often overlaps with regions of transcriptomic and epigenomic variability, and is influenced in part by genetic variation that may contribute to disease risk.

## RESULTS

### Mapping 3D chromatin conformation across individuals

To generate maps of 3D chromatin conformation suitable for comparison across individuals, we performed “dilution” Hi-C on LCLs derived from 13 Yoruban individuals (including one trio), one Puerto Rican trio, and one Han Chinese trio (19 individuals total; Supplemental Table 1). We also include published Hi-C data from one European LCL (GM12878) generated previously by our group using the same protocol^43^, for a total of 20 individuals from four different populations. Many of these same LCLs have been used in previous genomic studies^38,40,42^, allowing us to leverage multiple transcriptomic and epigenomic datasets in our analysis below (Supplemental Table 2). Importantly, 18 of these individuals have had their genetic variation cataloged by the 1000 Genomes Consortium^35,44^ (Supplemental Table 1), which allowed us to examine the influence of genetic variation on 3D chromatin conformation. Two replicates of Hi-C were performed on each LCL, with each replicate performed on cells grown independently in culture for at least two passages (Supplemental Table 3).

All Hi-C data were processed using a uniform pipeline that incorporates the WASP approach^40,45^ to eliminate allelic mapping biases (see **methods** section 2a). For each sample, we mapped a series of well-established Hi-C-derived features including 40Kb resolution contact matrices, Directionality Index (DI)^16^, Insulation score (INS)^7^, and compartmentalization^13^ (Figure 1a; Supplemental Figure 1a-c). Compartmentalization is measured by the first Principal Component (PC1) of Hi-C contact matrices, and thus we use the acronym “PC1” below to refer to this measure of compartmentalization. We also identified regions known as Frequently Interacting Regions (FIREs)^19^ and their corresponding “FIRE scores”, which measure how frequently a given region interacts with its neighboring regions (15~200kb). The concept of FIRE is based on the observation that the frequency of contacts at this distance is not evenly distributed across the genome, but rather, tends to peak in regions showing epigenomic signatures of transcriptional and regulatory activity (Supplemental Figure 2). As we have shown previously^19,20^, FIRE regions often overlap putative enhancer elements (Supplemental Figure 1d-e). We did not call “chromatin loops” in this study because our data was not of sufficient resolution, but we use a set of loops called previously in the LCL GM12878^14^ to examine variation in loop strength among the LCLs in our study. Aggregate analysis shows that these published LCL loops are generally reproduced in our data (Supplemental Figure 3).

**Figure 1.**
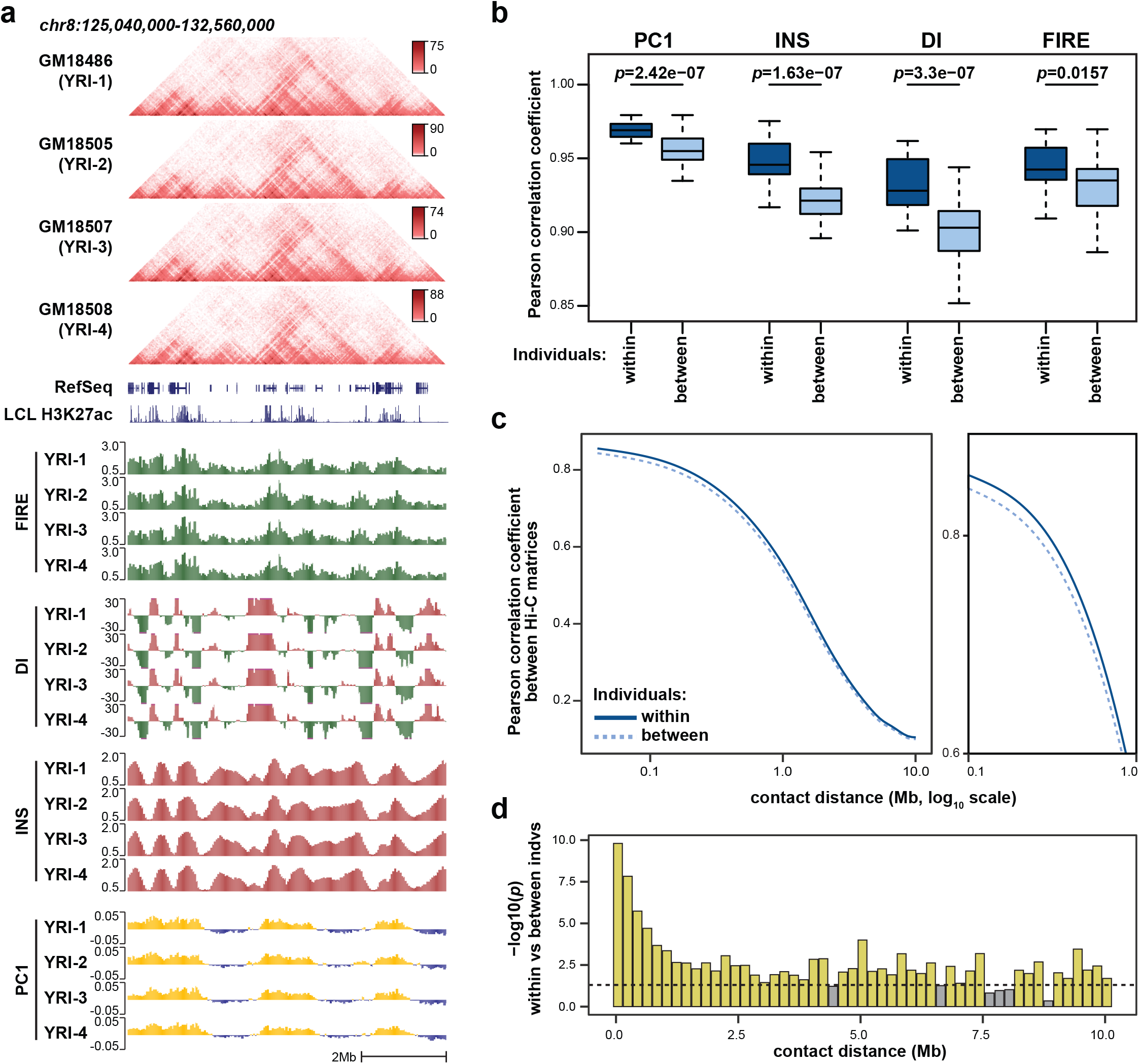
Biological variability in multiple aspects of 3D chromatin. (a) Browser view to illustrate the Hi-C-derived molecular phenotypes examined here: contact matrices, FIRE, DI, INS, and PC1 (chr8:125,040,000-132,560,000; hg19). Only 4 individuals shown here for illustrative purposes. The full set of individuals is shown in Supplemental Figure 1. (b) Boxplots show correlation between biological replicates from the same cell line (Individuals = “within”, N = 20), and between replicates from different cell lines (Individuals = “between”, N = 760). Statistical significance calculated by two-sided Wilcoxon rank sum test. See Supplemental Figure 4a for schematic of shuffling strategy. These and all boxplots in this manuscript show median as a horizontal line, interquartile range (IQR) as a box, and the most extreme value within 1.5*IQR or −1.5*IQR as whiskers extending above or below the box, respectively. (c) The Pearson correlation coefficient between quantile normalized Hi-C matrix replicates from the same cell line or different cell lines is plotted as a function of genomic distance between anchor bins. Distances between 0.1-1Mb are highlighted in the magnified sub-panel to the right. (d) Significance of the difference between the “within” and “between” values in (c) was calculated at multiple points along the distance-correlation curve by two-sided Wilcoxon rank sum test. Note that the scale of contact distance here is linear. Yellow bars indicate significance exceeding a nominal p-value of 0.05 (dotted line).

### 3D chromatin conformation variations between individuals

After uniformly processing all Hi-C data (see **methods** section 2), we compared chromatin conformation across LCLs at the level of contact matrices and multiple derived features (PC1, DI, INS, and FIRE). From a genome-wide perspective, each of these 3D chromatin features shows a signature consistent with reproducible inter-individual variation whereby replicates from the same individual (i.e. same LCL) are more highly correlated than datasets from different individuals (PC1 *p*=2.4e-7, INS *p*=1.6e-7, DI *p*=3.3e-7, FIRE *p*=0.0157 by Wilcoxon rank sum test; Figure 1b-d, Supplemental Figure 4a-f). The Hi-C data also cluster by population (Supplemental Figure 4f-g) consistent with an influence from genetic background, but we note that this population-level clustering can be caused by other factors such as batch of sample acquisition^46^.

Despite generally high correlations of Hi-C data across individuals, we frequently observed regions where 3D chromatin conformation varies reproducibly between individuals (example shown in Figure 2a, Supplemental Figure 5a). To more systematically identify regions of variable 3D chromatin conformation, we used the “*limma*” package^47^ to identify regions where variation between individuals was more significant than variation between two replicates from the same individual. We applied this approach to DI, INS, FIRE, and PC1. For each metric, we first defined a set of testable 40kb bins across the genome by filtering out bins with low levels of signal across all individuals or near structural variants that can appear as aberrations in Hi-C maps^48^ (see **methods** section 4a). We then applied a False Discovery Rate (FDR) threshold of 0.1 and merged neighboring variable bins, resulting in the identification of 2,318 variable DI regions, 2,485 variable INS regions, 1,996 variable FIRE regions, and 7,732 variable PC1 regions (Figure 2b, Supplemental Table 4, Supplemental Figure 5b). We note that there is strong overlap between the variable DI, INS, FIRE, and PC1 regions detected across all 20 LCLs and those detected using only the 11 unrelated YRI LCLs, which suggests that potential confounding effects of variation between different populations are not driving the identification of these variable regions (Supplemental Figure 5c). Although each metric has a unique set of testable bins, we found significant enrichment for bins that are variable in more than one metric (Figure 2c, Supplemental Figure 5d-e), indicating that the same regions often vary across multiple features of 3D chromatin conformation.

**Figure 2.**
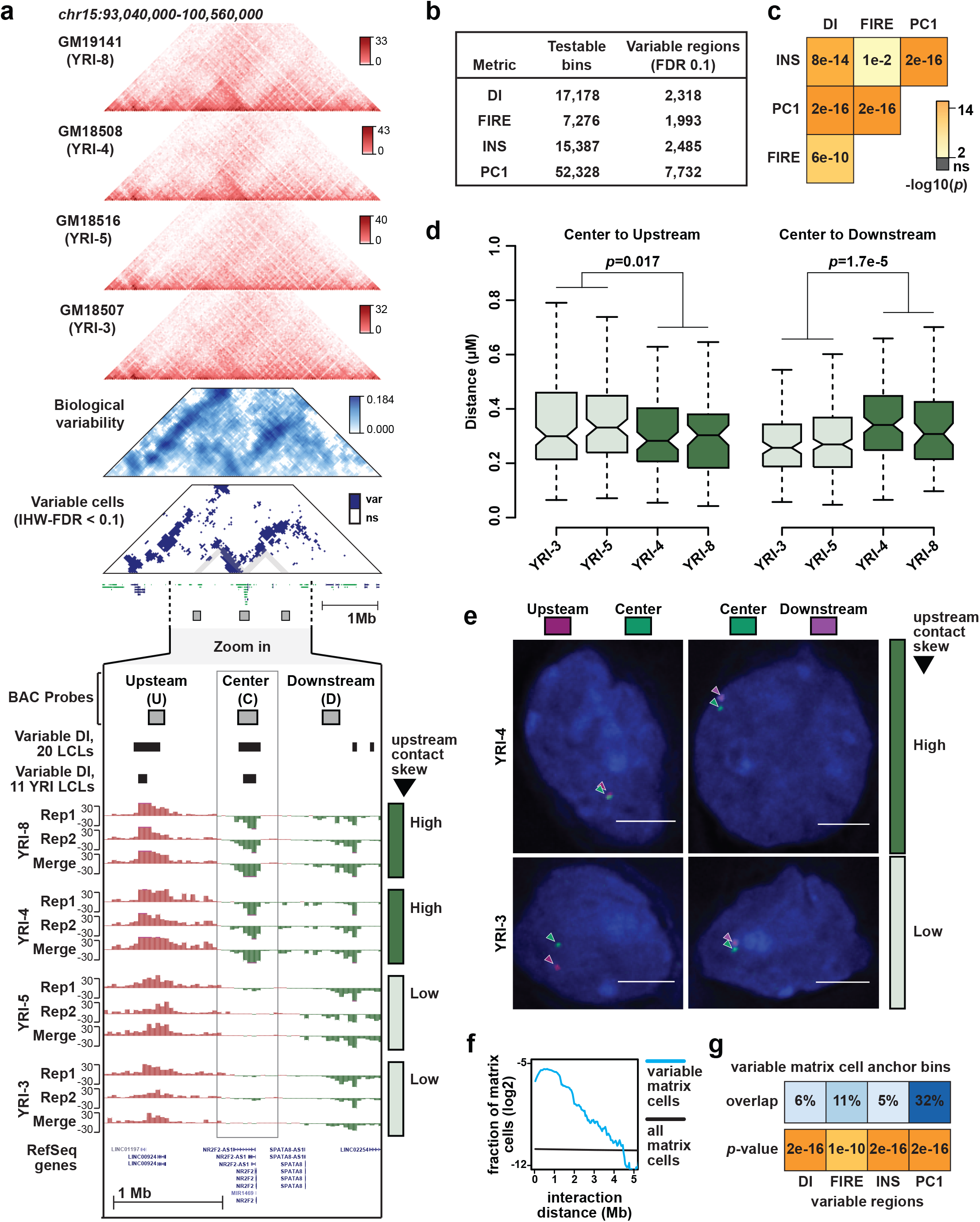
Variable regions of 3D chromatin conformation. (a) Example of a variable region (chr15:93,040,000-100,560,000; hg19). Triangular heatmaps from top to bottom: Four Hi-C contact heatmaps in red from individuals showing variable 3D chromatin architecture, a heatmap in blue showing the degree of variation measured across LCLs, and a heatmap in blue showing variable cells in the matrix at IHW-FDR < 0.1 (var=variable, ns=not significant). Standard tracks from top to bottom, and zoomed in more closely on the region of interest (chr15:95,482,152-98,025,591; hg19): BAC probes used for FISH experiment in panels 2d-e, variable DI regions called using all 20 LCLs, variable DI regions called using just 11 YRI LCLs, twelve DI tracks from four different individuals. For each individual, DI tracks are shown from two biological replicates and from Hi-C data merged across both replicates. Note that two individuals have strong upstream contact skew in the boxed regions (YRI-4, YRI-8), while the other two individuals have weak or no upstream contact skew in that region (YRI-3, YRI-5). (b) The number of testable bins and significantly variable regions for each 3D chromatin phenotype examined here. (c) Significance of pairwise overlap between different sets of variable regions. P values calculated by chi square test. Additional details in methods and Supplemental Figures 5 and 6. (d) Boxplots showing the distance between indicated probe sets in four different LCLs. Probe labels same as in panel (a). P values calculated by two-sided Wilcoxon rank sum test. Number of nuclei measured for each LCL and probe pair, from left to right, are: 140, 91, 111, 70, 128, 124, 219, 70. (e) Representative images of nuclei corresponding to panel (d). (f) Blue line shows the fraction of variable matrix cells distributed across a range of interaction distances. Black shows the fraction of all matrix cells distributed across the same range of interaction distances. (g) Top panel shows the percentage of variable matrix cell anchor bins that overlap variable DI, FIRE, INS, or PC1 regions, respectably. The shade of blue is scaled with overlap percentage. Bottom panel shows the statistical significance of these overlaps as calculated by chi square test, and plotted with same color scale as (c).

We next used Fluorescent In Situ Hybridization (FISH) to examine whether variable regions detected by Hi-C are consistent with distance measurements from imaging data (Figure 2d-e). Focusing on a variable DI region on chromosome 15 (chr15:96720000-96920000; hg19), we performed FISH in LCLs from four individuals with different levels of DI at the variable region being evaluated (YRI-3, YRI-4, YRI-5, YRI-8). We used three BAC probes that hybridize respectively to the variable DI region (“center”, probe covers chr15:96715965-96898793), a region approximately 668Kb upstream (“upstream”, probe covers chr15:95897555-96047720), or a region approximately 590Kb downstream (“downstream”, probe covers chr15:97488414-97648104). We found that distances between the center probe and these flanking probes vary significantly between individuals with strong upstream contact bias as measured by DI (YRI-4, YRI-8) and individuals without this upstream contact bias (YRI-3, YRI-5)(Figure 2d-e, center-upstream distance *p*=0.017, center-downstream distance *p*=1.7e-5 by Wilcoxon rank sum test). Moreover, we found that the center probe is closer to the upstream than the downstream probe in the two individuals with strong upstream DI signal at the central variable DI region (*p*=3.2e-3 for YRI-3, *p*=1.5e-4 for YRI-5 by Wilcoxon rank sum test). However, this trend is reversed in individuals without upstream DI signal where the center probe is now closer to the downstream probe (*p*=0.021 for YRI-4, *p*=0.1 for YRI-8 by Wilcoxon rank sum test) (Supplemental Figure 6a).

We also sought to identify variable entries in the Hi-C contact matrix itself (“matrix cells”). To facilitate this search, we used a method called Bandwise Normalization and Batch Correction (BNBC) that we recently developed to normalize Hi-C data across individuals (Fletez-Brant et al. Pre-print: https://doi.org/10.1101/214361). BNBC takes contact distance into account as a co-variate because batch effects in Hi-C data can be distance-dependent. To identify variable matrix cells, we performed a variance decomposition on Hi-C contact matrix cells which exhibited statistically-significant variability between individuals, resulting in a measure of biological variability for each bin in the contact matrix (see example in Figure 2a and Supplemental Figure 5a). To identify matrix cells with significant levels of biological variability, we estimated FDR using the IHW framework^49^ to include the distance between anchor bins as an informative covariate. At an FDR threshold of 0.1, we identified 115,817 matrix cells showing significant variability between samples (Supplemental Table 5). These variable bins are heavily skewed toward shorter contact distances (Figure 2f, Supplemental Figure 6b), likely due in part to higher read counts and thus increased power at these distances. We observed that the anchor regions of variable matrix cells overlap with variable regions of DI, INS, FIRE, and PC1 more often than would be expected by chance (Figure 2g; Supplemental Figure 6c). We also observed that variable matrix cells tend to occur in groups (Figures 2a, 3a), suggesting that variation in 3D chromatin conformation often affects more than one adjacent genomic window.

**Figure 3.**
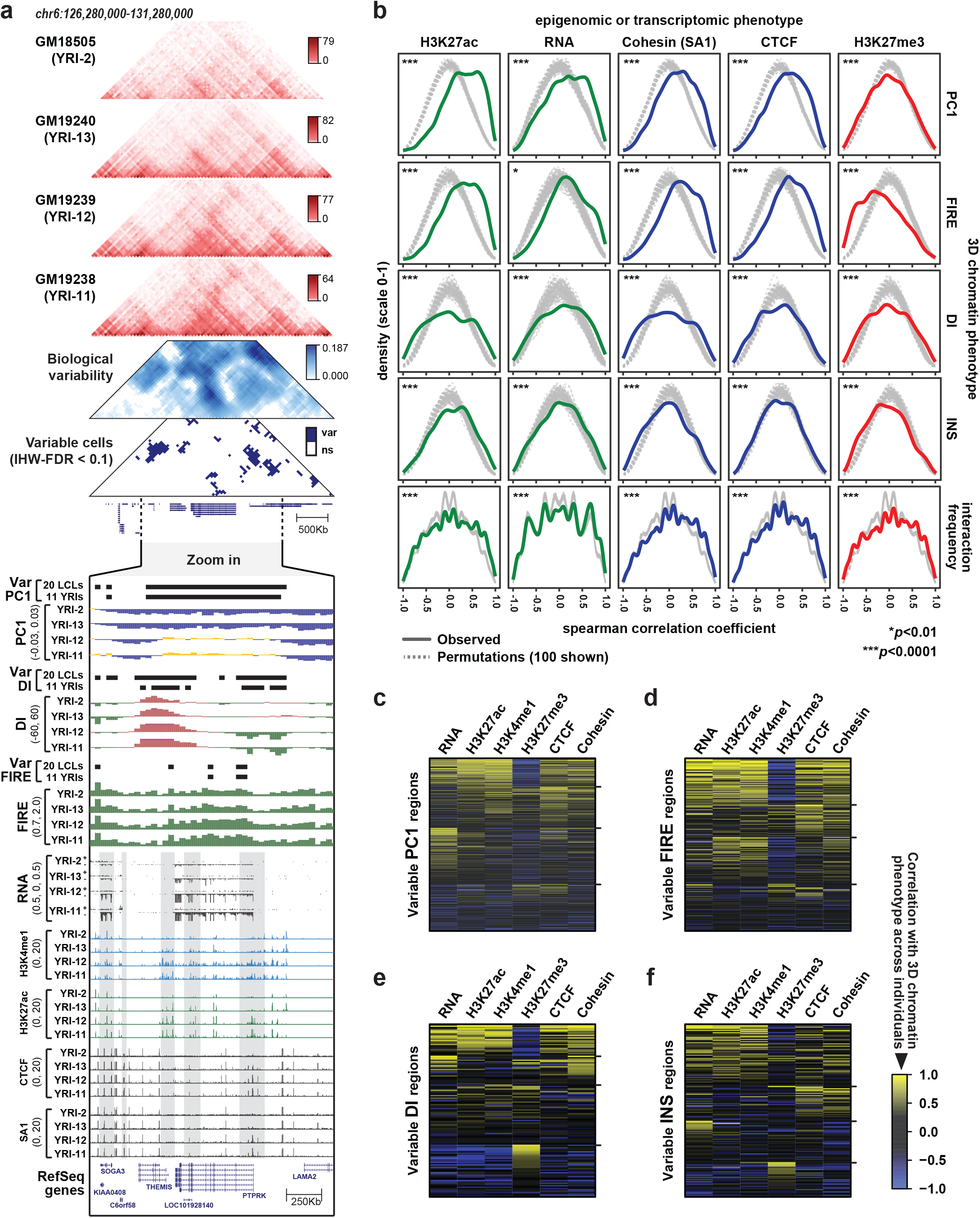
Coordinated variation of the 3D genome, epigenome, and transcriptome. (a) Example of a variable region where 3D chromatin phenotypes are correlated with epigenomic and transcriptomic phenotypes (chr6:126,280,000-131,280,000; hg19). Six triangular heatmaps from top to bottom: Hi-C contact heatmaps from four individuals, variability matrix, and variable cells in the matrix (var=variable, ns=not significant). Standard tracks below show 3D chromatin, epigenomic, and transcriptomic properties from four individuals in zoomed in region (chr6:127,680,918-129,416,097; hg19). All ChIP-seq and RNA-seq data in the figure from Kasowski et al, 2013^42^. (b) Density plots show the distribution of Spearman correlation coefficients at variable regions (see Figure 2b for numbers) between the epigenomic or transcriptomic phenotype indicated in the top margin of panel and the 3D chromatin phenotype indicated in the right margin of panel. Grey lines show the distributions from 100 random permutations (selected randomly from the 10,000 permutations performed) in which the sample labels were shuffled (see Supplemental Figure 7b). ***p<0.0001 by permutation test as described in methods section 4c, which applies to all observations in this panel except RNA-seq at INS regions (*p*=0.0018) and RNA-seq at FIRE regions (*p*=0.0096). (c) Heatmap showing Spearman correlation coefficients between PC1 and multiple epigenomic/transcriptomic phenotypes, arranged by k-means clustering (k=4). Tick marks to the right show boundaries between clusters. Each row (N=518) is one variable PC1 region, limited to the subset of variable PC1 regions that contain RNA-seq signal and at least one peak in at least one individual for each ChIP-seq target included here (H3K27ac, H3K4me1, H3K27me3, CTCF, Cohesin). (d) Similar to (c), showing correlations with FIRE at N=132 variable FIRE regions. (e) Similar to (c), showing correlations with DI N=265 variable DI regions. (f) Similar to (c), showing correlations with INS at N=154 variable INS regions.

### Coordinated variation of the 3D genome, epigenome, and transcriptome

To investigate the relationship between variation in 3D chromatin conformation and gene regulation, we analyzed multiple published datasets including RNA-seq, ChIP-seq, and DNase-seq data generated from some of the same LCLs in our study (Supplemental Table 2). Strikingly, for all external datasets examined here, we see an enrichment for regions at which 3D chromatin conformation across individuals is correlated with measures of genome activity in the same 40Kb bin (see example in Figure 3a and Supplemental Figure 7a). To assign a level of statistical significance to these observations, we approximated the null distribution by randomly permuting the sample labels of external datasets, thus disrupting the link between Hi-C and ChIP/RNA/DNase-seq data from the same individual, but not changing the underlying data structure (see schematic in Supplemental Figure 7b). We used these permutations to calculate the bootstrap p-values in Figure 3b. Among variable PC1 regions, we observed a significant enrichment for regions at which PC1 values across individuals are positively correlated with histone modifications indicative of transcriptional activity including H3K27ac (bootstrap P<0.001), H3K4me1, and H3K4me3 (but notably less so with H3K27me3, which is marker of transcriptional repression) (bootstrap P<0.001 for all histone modifications, Figure 3b). The correlations between PC1 and marks of transcriptional activity occur in the expected direction – i.e. higher PC1 values are associated with higher gene expression and more active histone modifications. Similar correlations were apparent in two distinct sets of ChIP-seq data generated by different groups^40,42^, and observed whether we use variable regions identified across all 20 LCLs or only across the 11 unrelated YRI LCLs (Supplemental Figure 7c).

The relationship between variation in 3D chromatin conformation and underlying genome activity extends beyond A/B compartmentalization. At variable FIRE regions, we found an abundance of regions where FIRE score is positively correlated with marks of cis-regulatory activity including H3K27ac and H3K4me1 (Bootstrap P<0.001; Figure 3b, Supplemental Figure 9a), consistent with the previously reported relationship between FIREs and cis-regulatory activity^19,20^. DI and INS values at variable regions tend to be correlated histone modification levels as well as CTCF and Cohesin subunit SA1 binding (Bootstrap P<0.001; Figure 3b, Supplemental Figure 8a-b), which are known to influence these 3D chromatin features^16,50,51^. For INS, the relationship is directional such that higher CTCF/Cohesin binding corresponds to more contact insulation (i.e. lower INS score). However, at variable DI regions the correlations are not as clearly directional, reflecting current understanding that the direction of DI (i.e. upstream vs downstream contact bias) is arbitrary relative to strength of CTCF/Cohesin binding. We performed similar analysis on variable cells in the contact matrix, and found that the interaction frequency in these matrix cells across individuals tends to be correlated with epigenetic or transcriptional properties of one or both corresponding “anchor” bins (Bootstrap P<0.001; Figure 3b, Supplemental Figure 9b). Importantly, for all types of variable regions examined here we found correlation with RNA-seq signal, indicating that at least at some regions, variation in 3D chromatin features accompanies variation in gene expression.

We examined further whether 3D chromatin conformation at a given variable region tends to be correlated with only one epigenomic property, or with several properties simultaneously. We found that PC1, FIRE, INS, and DI values across individuals are often correlated with multiple features of active regions (e.g. H3K27ac, H3K4me1, RNA), and anti-correlated with the repressive H3K27me3 histone modification (Figure 3c,d). For DI, where direction is not as clearly linked to magnitude of gene regulatory activity, we note a larger set of regions with anti-correlation to features of active regions (e.g. H3K27ac, H3K4me1, RNA) and positive correlation with H3K27me3 (Figure 3e,f). These results demonstrate that variation in 3D chromatin conformation is often accompanied by variation in transcriptional and regulatory activity of the same region. Moreover, the correlations between multiple molecular phenotypes at the same region suggest that shared mechanism(s) underlie variation in these phenotypes across individuals.

### Genetic loci influencing 3D chromatin conformation

To examine genetic influence on 3D chromatin conformation we first considered genetic variants overlapping CTCF motifs at chromatin loop anchors^14^, because disruption of these CTCF motifs by genome engineering has been shown to alter chromatin looping^23^. Focusing on SNPs at variation-intolerant positions in anchor CTCF motifs (“anchor disrupting SNPs”, at sequence weight matrix positions where a single base has a probability of >0.75, Figure 4a), we observed a significant linear relationship between SNP genotype and the strength of corresponding loops (*p*=7.6e-5 by linear regression; Figure 4b,c). We also examined whether individuals heterozygous for anchor disrupting SNPs showed allelic imbalance in loop strength. To facilitate this analysis, we used the HaploSeq^43^ method to generate chromosome-span haplotype blocks for each LCL (Supplemental Table 6). Although few Hi-C read pairs overlap a SNP allowing haplotype assignment (mean 7.89% of usable reads per LCL), we do observe that the haplotype bearing the stronger motif allele tends to show more reads connecting the corresponding loop anchors (*p*=5.9e-4 by one-sided t-test of mean > 0.5; Figure 4d). Our observation that CTCF motif SNPs can modulate 3D chromatin conformation is consistent with similar findings reported from ChIA-PET data^52^, and a recent report of haplotype-associate chromatin loop published while this manuscript was in preparation^27^.

**Figure 4.**
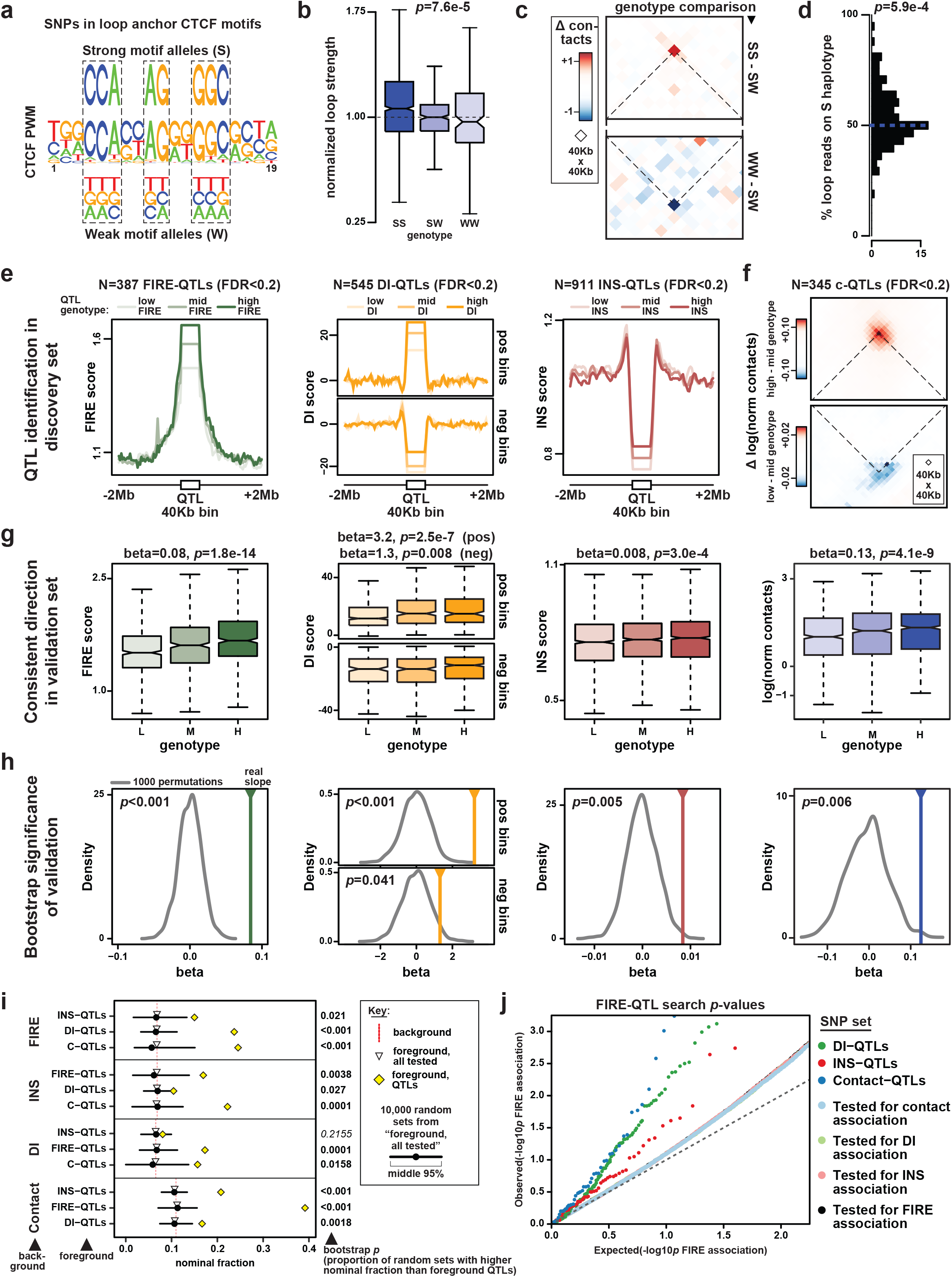
A genetic contribution to variations in 3D chromatin conformation. (a) A graphic representation of the CTCF Position Weight Matrix (PWM) is shown. Eight positions boxed by dashed lines have probability >0.75 for a single base. We refer to SNPs at these positions as “motif disrupting SNPs”. Alleles matching the consensus base in the motif are labeled “strong motif alleles (S)”, and alleles matching any other base are labeled “weak motif alleles (W)”. (b) Boxplot shows the distribution of interaction frequencies at loops with exactly one anchor containing a CTCF motif disrupting SNP (N=138), separated according to genotype. For each SNP, loop strengths are normalized to the mean value of the heterozygous genotype (WS). There is significant linear relationship between normalized loop strength and genotype by linear regression (*p*=7.6e-5). (c) Aggregate contact map shows the average difference in interaction frequency per loop between SS and SW genotypes (top; N=117 SNPs), and between SW and WW genotypes (bottom; N=31 SNPs). The cross point of dotted lines indicates the 40Kb bin containing the loop being evaluated. (d) Histogram shows the allelic imbalance in reads connecting loop anchors on the S vs W haplotypes in WS heterozygotes (N=135 loops). The mean percentage of reads on the S haplotypes is significantly larger than 0.5 (*p*= 5.9e-4 by onesided t-test). (e) Line plots show the genotype-dependent signal of FIRE-QTL, INS-QTL and DI-QTL using 11 independent YRI individuals. Each plot show the indicated phenotype as lines with light color, medium color and dark color representing average signal across LCLs with the low signal genotype, medium signal genotype, and high signal genotype, respectively. For DI-QTL, we split all 40Kb QTL bin into two groups, based on either upstream DI bias (upper panel) or downstream DI bias (bottom panel). (f) For C-QTLs, an aggregate contact plot analogous to panel c is used to show the average difference in BNBC corrected interaction frequency (“Δ log(norm contacts)”) between the high and medium contact genotypes (top; N=138 interactions), and between the genotypes medium and low genotypes (bottom; N=94 interactions). The cross point of dotted lines indicates the 40Kb test bin in question. (g) Boxplots show the genotype-dependent signal at QTLs using additional 6 individuals as a validation set. In each boxplot, three boxes with light color, median color and dark color represent the average signal in the 40Kb QTL bin from individuals with the low signal genotype, medium signal genotype, and high signal genotype, respectively. (h) Results of permutation test to evaluate the statistical significance of results in (g). The solid vertical lines show the estimated linear regression slope values obtained from the validation set (N=6 individuals). The grey curves show the distributions of slope values obtained from 1,000 random permutations. Corresponding bootstrap *p* values indicated in the upper left corner of each subpanel. (i) Line plot shows the fraction of foreground SNPs with nominal significance in the background association study (“nominal fraction”). Red dashed lines show the values for all SNPs in a given background set, yellow diamonds show values for SNPs in a given foreground set, and open triangles show values for SNPs tested in the foreground QTL search. Black circles and lines indicate the median and middle 95% range, respectively, of 10,000 permutations in which SNPs were selected from the “foreground, tested” set. The number to the right of each line indicates the fraction of permutations with a value higher than observed for “foreground, QTL” set. (j) QQ Plot shows FIRE-QTL search results, including all SNPs tested for FIRE association (black points, N=128,137), and several subsets as follows: DI-QTL tested (light green, N=46,784), INS-QTLs tested (light red; N=6,238), C-QTL tested (light blue; N=69,847), DI-QTLs (dark green, N=152), INS-QTLs (dark red, N=60), C-QTLs (dark blue, N=53).

Motivated by these preliminary observations of genetic effects on 3D chromatin conformation, we next searched directly for QTLs associated with Hi-C derived features of 3D chromatin conformation. Power calculations indicated that, despite limited sample size, we were moderately powered to find QTLs with strong effect sizes using a linear mixed effect model (LMM) approach that takes advantage of the Hi-C replicates for each LCL (Supplemental Table 7). Thus, we conducted a targeted search for QTLs associated with variation in FIRE, DI, INS, and contact frequency. We did not include PC1 in the QTL search because we reasoned that individual genetic variants would be more likely to have detectable effects on local chromatin conformation rather than large-scale features like compartmentalization. For this same reason, we used modified versions of DI and INS scores for the QTL search calculated with a window size of 200Kb upstream and downstream of the target bin, rather than the standard 2Mb window size for DI^16^ or 480Kb for INS^47^. We also limited our QTL searches to the 11 unrelated YRI individuals in our study (referred to below as the “discovery set”) to mitigate potential confounding differences between populations.

For each 3D genome phenotype under study we identified a list of testable bins that showed appreciable levels of signal in at least one individual in our discovery set (see **methods** section 7 for full description of test bin and SNP selection). We also identified a set of test SNPs that includes at most one tag SNP among those in perfect LD in each 40Kb bin. Response variables (i.e. 3D chromatin phenotype values) were quantile normalized across the discovery set. For each testable bin, we measured the association of the given 3D chromatin phenotype with all test SNPs in that bin. In cases where multiple SNPs in the same bin were significantly associated with the phenotype we selected only the most significantly associated SNP per bin for our final QTL list. Ultimately, at an FDR of 0.2, we identified 387 FIRE-QTLs (i.e. testable bins in which FIRE score is associated with at least one SNPs in that bin; comprising 6.6% of tested bins), 545 DI-QTLs (4.2% of tested bins), and 911 INS-QTLs (12.0% of tested bins)(Figure 4e, Supplemental Figure 10a, Supplemental Table 8). For analysis of DI-QTLs, we separated the testable bins into those with upstream bias and those with downstream bias (see **methods** section 7d), because we observed a Simpson’s paradox when we analyzed the genotype trend at all DI-QTL regions together (Supplemental figure 10b).

We also searched for QTLs associated directly with interaction frequency in individual contact matrix cells using an LMM approach like that described above for FIRE, DI, and INS. The large number of cells in a Hi-C contact matrix, together with limited sample size, made a true genome-wide QTL search unfeasible. However, power calculations indicated that if we limited our QTL search to a subset of cells in the matrix we could have moderate power to detect strong genetic signals (Supplemental Table 7). Thus, we limited our QTL search for contact matrix QTLs (“C-QTLs”) to matrix cells that showed significant biological variability in our samples, as described above. We tested for association in our discovery set between the BNBC-normalized interaction frequency in these variable matrix cells and the genotype of test SNPs in either of the two anchor bins. We selected at most one QTL SNP per matrix cell, using association p-value to prioritize, finally yielding 345 C-QTL SNPs associated with 463 matrix cells at an IHW-FDR threshold of 0.2 (Figure 4f, Supplemental Table 8).

To evaluate the reproducibility of each of these QTLs sets (FIRE-QTLs, DI-QTLs, INS-QTLs, and C-QTLs), we examined Hi-C data from 6 individuals who were not included in our discovery set (we refer to these 6 individuals our “validation set”; Supplemental Table 1). These individuals represent four different populations (CEU, PUR, CHS, YRI), and they include a child of two individuals in the discovery set (YRI-13/NA19240 is child of YRI-11/NA19238 and YRI-12/NA19239). In each case, we find a significant linear relationship in the validation set between QTL genotype and the corresponding 3D chromatin phenotype (*p*=1.8e-14 for FIRE-QTLs, *p*=2.5e-7 for DI-QTLs at positive DI bins, *p*=0.008 for DI-QTLs at negative DI bins, *p*=3e-4 for INS-QTLs, *p*=4.1e-9 for C-QTLs; Figure 4g). To provide an additional and more stringent estimate of the significance of these observations, we performed permutations by randomly selecting sets of test SNPs and measuring the linear relationship between genotype and phenotype in the validation set. In all cases, the observed relationship was also significant by this more conservative bootstrap approach (p<0.001 for FIRE-QTLs, *p*<0.001 for DI-QTLs at positive DI bins, *p*=0.041 for DI-QTLs at negative DI bins, *p*=0.005 for INS-QTLs, *p*=0.006 for C-QTLs; Figure 4h).

There is little direct overlap between our different QTL sets (Supplemental figure 10c), likely due to limited power and the fact that the testable bins were different for each metric. However, we observed genotype-dependent INS score at FIRE-QTLs and C-QTLs, and genotype-dependent FIRE score at INS-QTLs and DI-QTLs (Supplemental Figure 10d), which suggested that overlapping signal between different types of 3D chromatin QTLs in present below the level of test-wide significance. To more rigorously assess overlapping signal between our QTL sets we examined shared association below the threshold of multiple test correction, inspired by similar approaches reported elsewhere^53^. Our underlying hypothesis is that genetic association studies of two different phenotypes “*X*” and “*Y*” with overlapping (or partially overlapping) genetic architecture may have few direct overlaps between significant hits due to limited power or differing study designs, but the shared signal should become apparent when the full range of association results are considered. To quantify this, we calculated the fraction of QTLs for a given phenotype *X* that exceed a nominal level of significance (*p* < 0.05) when tested for association with a different phenotype *Y*. We refer to this value as the “nominal fraction” below and in figure 4i. To test whether the nominal fraction of *X*-QTLs was significantly higher than would be expected by chance, we approximated the null distribution by calculating nominal fractions for 10,000 sets of SNPs selected randomly from among all *X* test SNPs. In almost all pairwise comparisons between 3D chromatin QTL types examined here, we find that the observed nominal fractions are significantly higher than would be expected in the absence of shared genetic architecture (Figure 4i,j).

### Contribution of 3D chromatin QTLs to molecular phenotypes and disease risk

Given the correlation observed between 3D chromatin variation and epigenome variation, we next investigated whether 3D chromatin QTLs could modulate both the epigenome and 3D genome. Here, we made use of published ChIP-seq data for histone modifications (H3K4me1, H3K4me3, H3K27ac) in a large set of 65 YRI LCLs^39^, DNase-seq data from 59 YRI LCLs^38^, and CTCF ChIP-seq data from 15 CEU LCLs^54^. Notably, most individuals in these datasets were not included in our QTL discovery or validation sets (54/65 for histone modification ChIP-seq, 48/59 for DNase-seq, 15/15 for CTCF ChIP-seq). In many cases, we found a significant linear relationship between 3D chromatin QTL genotypes and these different epigenetic phenotypes (Figure 5a, Supplemental figure 11a). For example, at FIRE-QTLs, the high-FIRE allele is also associated with higher levels of active histone modifications and chromatin accessibility (Figure 5a). We note that although these associations are all significant by linear regression, only H3K27ac and H3K4me1 passed more conservative permutation testing in which the null distribution is approximated by selecting random SNPs from the full set of tested SNPs (Figure 5b). At C-QTLs, the high-contact alleles show higher levels of the enhancer-associated mark H3K4me1 in the two anchor bins that connect the corresponding matrix cell. Moreover, the nominal fraction of C-QTLs (i.e. fraction of c-QTLs with *p*<0.05) in a published set of H3K4me1-QTLs is significantly higher than expected in the absence of shared genetic association (*p*=6.9e-6 by chi square test, bootstrap *p*=0.028; Supplemental Figure 11b,d). At INS-QTLs, the slope of these genotype-phenotype relationships is inverted such that higher levels of histone modifications and chromatin accessibility are associated with the low INS score allele (i.e. more contact insulation), although only the association with chromatin accessibility is significant by both linear regression and permutation test (*p*=1.6e-40 by linear regression, bootstrap *p*=0.023; Supplemental Figure 11b,d). The genotype-phenotype relationships observed at DI-QTLs are not as clear as for other metrics (Figure 5b, Supplemental figure 11a), but this is expected because increased histone modifications or chromatin accessibility can influence DI in either direction, potentially confounding this type of aggregate analysis. Anecdotally, we do observe examples of individual DI-QTLs where genotype appears to correlate with epigenomic phenotype (Figure 5c).

**Figure 5.**
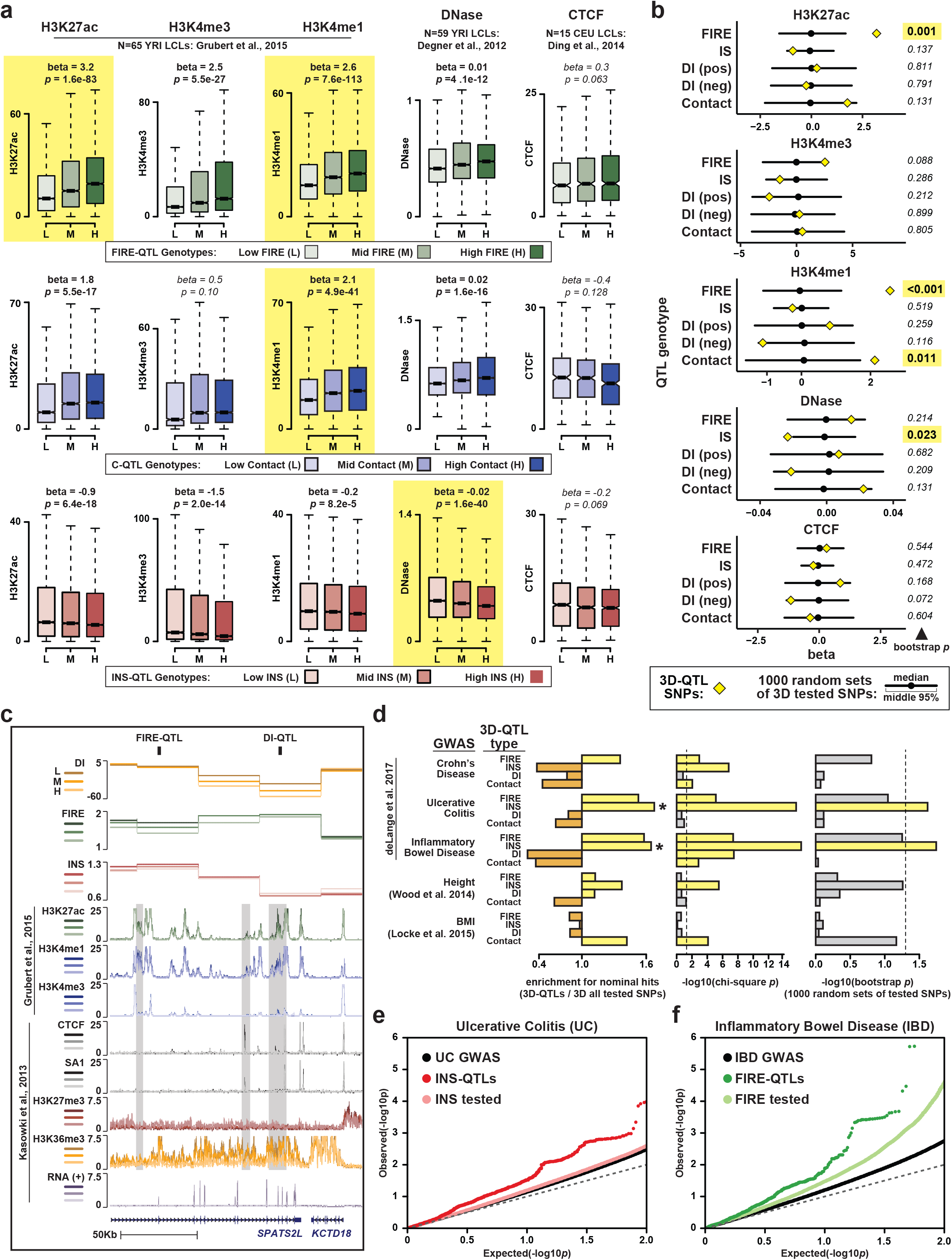
Contribution of 3D chromatin QTLs to other molecular and organismal phenotypes. (a) Boxplots show signal for epigenetic phenotypes separated by genotype at FIRE-QTLs (top row), C-QTLs (middle row), and INS-QTLs (bottom row). Epigenetic signals averaged across all peaks in 40Kb bin. Linear regression *p* and beta values shown above each plot. P-value <0.05 in bold, others in italics. Yellow boxes highlight relationships that are significant by linear regression and in permutation testing as shown in (b). (b) Line plots shows beta values of linear relationships between QTL genotypes as indicated to the left and epigenetic phenotype indicated above each subpanel. Yellow diamonds show values for the true QTLs sets as shown in (a) and Supplemental figure 11a. Black circles and lines indicate the median and middle 95% range, respectively, of 1,000 permutations in which SNPs were selected from the “foreground, tested” set. The number to the right of each line indicates the fraction of permutations with abs(beta) higher than observed for the true QTL set. Yellow boxes highlight values < 0.05. (c) Genome browser view (chr2:201,222,342-201,386,844; hg19) showing examples of a DI-QTL (chr2:201333312) and FIRE-QTL (chr2:201254049). All signals plotted as a function of DI-QTL genotype (L=Low DI, M=medium DI, H=High DI). Grey boxes highlight region where epigenetic signals stratify by DI-QTL genotype. (d) Left subpanel shows the enrichment values for 3D QTL SNPs with nominal significance in the indicated GWAS study calculated as follows: (fraction of indicated 3D QTL SNPs with nominal significance in the indicated GWAS) / (fraction of SNPs tested in the indicated 3D QTL search with nominal significance in the indicated GWAS). Asterisks mark values with P<0.05 by chi-square test (middle panel), and permutation test (right panel). Right panel shows the proportion of 1000 random subsets selected from the tested SNPs with enrichment values higher than the indicated true QTL set. Dotted lines mark *p*=0.05. (e) QQ plot shows the results of UC GWAS with all tested SNPs shows as black points, and two subsets as follows: SNPs also tested in our INS-QTL search (light red), and SNP called as INS-QTLs or in perfect LD with INS-QTLs in the same 40Kb bin (dark red). (f) QQ plot shows the results of IBD GWAS with all tested SNPs shows as black points, and two subsets as follows: SNPs also tested in our FIRE-QTL search (light green), and SNP called as FIRE-QTLs or in perfect LD with FIRE-QTLs in the same 40Kb bin (dark green).

Finally, we sought to examine whether 3D chromatin QTLs might contribute risk for complex diseases. There are 44 direct overlaps between our 3D chromatin QTLs (or SNPs in perfect LD in the same 40Kb bin) and NHGRI-EBI GWAS catalog^55^ (Supplemental Table 9). However, the significance of these direct overlaps is hard to assess given the differences between the populations and study designs in question. Thus, here again we examined overlaps below the level of genome-wide significance by looking at nominal fractions to assess shared signal between association studies. We compiled full summary statistics for large GWAS (>50,000 individuals) of the related immune-relevant phenotypes Crohn’s Disease (CD), Ulcerative Colitis (UC), and Inflammatory Bowel Disease (IBD)^56^, as well as studies of the non-immune phenotypes height^57^ and Body Mass Index (BMI)^58^. We observed striking enrichments for INS-QTLs among variants with nominal associations to UC and IBD risk (1.67- and 1.65-fold, respectively), and these enrichments are significant by both chi square and permutation tests (INS-QTL with UC chi square *p*=2.5e-16 and bootstrap *p*=0.024; INS-QTL with IBD chi square *p*=5.5e-17 and bootstrap *p*=0.018; Figure 5d,e). We also note a trend in which FIRE-QTLs show nominal association with UC and IBD (1.36- and 1.58-fold enrichment, respectively), although these observations fall just below the threshold of significance by the more stringent permutation test (FIRE-QTL with UC chi square *p*= 7.6e-6 and bootstrap *p*=0.090; FIRE-QTL with IBD chi square *p*= 4.2e-8 and bootstrap *p*=0.056; Figure 5d,f).

## DISCUSSION

Our results provide the first systematic characterization of how chromatin conformation varies between unrelated individuals at the population level, and as a consequence of genetic variation. The most important finding of our study is that genetic variation influences multiple features of 3D chromatin conformation, and does so to an extent that is detectable even with limited sample size and Hi-C resolution. To the best of our knowledge, this represents the first report of QTLs directly associated with 3D chromatin conformation. However, there are limitations to our QTL search that are important to note here. First, the small sample size means that our power to detect QTLs is limited, and in order to identify QTL sets that could be analyzed in aggregate we tolerated elevated type I error by using an FDR threshold of 0.2 (as done previously for molecular QTL studies with limited power^40^). Second, the limited resolution of our Hi-C data (40Kb) and extensive LD in our study population prevented us from identifying specific causal variant(s) for validation through genetic perturbation experiments. Nonetheless, we were able to validate the 3D chromatin QTL sets through aggregate analysis of Hi-C data from a small set of individuals who were not included in the QTL search, and with independently generated ChIP-seq and DNase-seq data from a larger set of individuals. Taken together, our results show that genetic variation influences several features of 3D chromatin conformation, which is an important step forward to evaluate the role of 3D chromatin conformation in mediating disease risk.

Another key finding of our study is that regions which vary in 3D chromatin conformation across individuals also tend to vary in measures of transcriptional and regulatory activity. This supports the existence of shared mechanisms that underlie variation in 3D chromatin conformation, transcription, and epigenomic properties. We suspect that no single mechanism or causal hierarchy applies to all regions of the genome with variation in one or more of these properties. However, in at least some cases, this shared mechanism is likely genetic. This raises the question of whether 3D chromatin QTLs are fundamentally the same as QTLs previously described for other molecular phenotypes (e.g. eQTLs, dsQTLs, histoneQTLs; collectively referred to below as “molQTLs”), or represent a separate set of QTLs not detectable with other methods. This question is difficult to answer in the present study for two main reasons: 1) Our power is limited and thus we cannot say with confidence that a given SNP is *not* a 3D chromatin QTL. Many molQTL studies also have limited power and are thus prone to type II error. 2) Our QTL searches, like most molQTL studies, are not truly genome-wide because subsets of testable regions and testable SNPs are preselected to focus the search space. These selection criteria can differ widely between studies, making direct QTL-to-QTL comparisons challenging. The observation of genotype dependent epigenetic signal at 3D chromatin QTLs suggest that at least some 3D chromatin QTLs could also be detected as other types of molQTLs if those studies had sufficient statistical power. However, the limited overlap between 3D chromatin QTLs and published molQTLs (even when considering SNPs with only a nominal level of significance) points to a lack of power in current studies, and suggests further that the QTLs with largest effects on 3D chromatin conformation are not necessarily the same as those with large effects on other molecular phenotypes, and vice versa. Therefore, it is likely that QTL studies directed toward different types of molecular phenotypes (including 3D chromatin features) are likely to be complimentary rather than redundant.

Future studies with higher resolution Hi-C data and larger sample sizes will be important to identify functional variants modulating 3D chromatin conformation, and to further dissect the mechanistic relationships between genetics, 3D chromatin conformation, and other molecular phenotypes. We anticipate that these studies will continue to reveal cases in which perturbation of 3D chromatin conformation is a molecular mechanism through which disease-associated genetic variants confer disease risk. The present study provides initial discoveries of genetic influence on 3D chromatin conformation and an analytical framework and that we believe will facilitate future efforts to unravel the molecular basis of genetic disease risk.

## AUTHOR CONTRIBUTIONS

Study was conceived and overseen by B.R., M.H., K.H., J.S., K.G., and D.U.G. Hi-C experiments performed by D.U.G. and A.S. Data analysis performed by Y.Q., M.H., K.F-B, (variable matrix cells, C-QTLs, related contact matrix analyses, power calculations), A.N., Y.L., J.C., and D.U.G. FISH experiments performed by T.L. Manuscript written by D.U.G., Y.Q., M.H., K.F-B., with input from all authors.

## METHODS

### 1. Hi-C data generation

Hi-C was performed as previously described^13^. We note that all Hi-C experiments were performed using a “dilution” HindIII protocol, rather than the newer “in situ” version of the protocol, for consistency because data generation began before the invention of in situ Hi-C. In addition, the resolution of 40kb used here for most analysis was determined primarily by sequencing depth rather than choice of a restriction enzyme. Thus, even if a 4-cutter like MboI had been used, the prohibitive cost of sequencing would have prevented us taking advantage of the additional possible resolution.

### 2. Hi-C data processing

#### 2.a. Alignment with WASP

Read ends were aligned to the hg19 reference genome using BWA-MEM^63^ v0.7.8 as single-end reads with the following parameters: -L 13,13. We used the WASP pipeline^40,45^ to control for potential allelic mapping biases, which some modifications to account for unique aspects of Hi-C data. BWA-MEM can produce split alignments where different parts of a read are aligned to different parts of the genome. This is critical for Hi-C data, because a read can span a Hi-C ligation junction between two interacting fragments. In the case of a split alignment, BWA-MEM will mark the higher-scoring alignment as the primary alignment. For Hi-C data this is not ideal – we want the five-prime-most alignment (before the ligation junction) to be the primary alignment. To account for this, we further processed the alignments from BWA-MEM to select the five-prime-most alignment in cases where one read was split. Reads without an alignment to the five-prime end of the read were filtered out, as were alignments with low mapping quality (<10). The WASP pipeline was then used to generate alternative reads by flipping the allele in reads overlapping SNPs, and these reads were then realigned using the same pipeline. As input to WASP, we included all SNPs and indels present in the PUR individuals in our set (HG00731, 732, 733), CHS individuals in 1000 genomes (we included all CHS to account for the fact that no 1000 genomes genotype calls were available for HG00514), YRI individuals in 1000 genomes (we included all YRI individuals to account for the fact that no 1000 genomes genotype calls were available for GM19193), and the H1 cell line^21^ (to facilitate uniform processing and comparisons between LCLs and H1-derived datasets). After alignment of the alternative reads, alignment of the original reads and alternative reads were compared by WASP, and only the original reads for which all alternative reads aligned at the same location with same CIGAR string were kept. Reads overlapping indels were removed. Reads were then re-paired, and only pairs in which both reads survived this filtering were kept. PCR duplicates were removed using Picard tool (http://broadinstitute.github.io/picard/) with default parameters. To ensure that our adapted WASP pipeline removed allelic mapping biases effectively, we simulated all possible 100bp single end reads spanning SNPs in our LCLs and aligned them back to the genome using our adapted WASP pipeline. We found no SNPs which depart from 50/50 mapping ration between reference and alternative allele in these simulations.

We also took steps to remove any potential artifacts due to HindIII polymorphisms. Hi-C data was obtained by cutting the genome with HindIII, so we reasoned that SNPs or indels that disrupt existing HindIII sites or create novel HindIII sites could lead to differential cutting of two alleles and thus the appearance of differential contact frequency. To mitigate these potential artifacts, we identified all HindIII sites that would be disrupted or created by genetic variants present in our samples, and removed all reads within 1Kb of these polymorphisms in all individuals.

#### 2.b. Contact Matrix Calculations

Matrices were generated and normalized as previously described^21,64^. Briefly, intra-chromosomal read pairs were divided into 40Kb bin pairs based on five prime positions. The number of read pairs connecting each pair of 40Kb bins were tallied to produce contact matrices for each chromosome. Raw counts in the contact matrices were then normalized using HiCNorm^64^ to correct for known sources of bias in Hi-C contact matrices (GC content, mappability, fragment length). Bins that are unmappable (effective fragment length, GC content or mappability is 0) were assigned NA values. These normalized matrices were further quantile normalized across samples to account for differing read depths and mitigate potential batch effects. One such quantile normalized matrix was generated for each chromosome in each replicate, as well as in each sample (replicates pooled together). We eliminated chromosomes X and Y from all downstream analyses due to the gender differences between our samples.

#### 2.c. PC1 Score

PC1 scores were computed using methods defined previously^13^. Briefly, quantile normalized matrices for each chromosome were transformed to Observed/Expected (O/E) matrices by dividing each entry in the matrix by the expected contact frequency between regions in that matrix at a given genomic distance. For a given matrix, the expected contact frequencies were computed by averaging contact frequencies at the same distance in that each matrix. The O/E matrices were further transformed to Pearson correlation matrices by the “cor” function in R and eigen vectors (principal components) were computed using the “cov” function in R. Generally, the first eigenvector (“PC1”) reflects A/B compartmentalization. However, for some chromosomes we have seen that the second eigenvector sometimes reflects compartmentalization, while the first eigenvector reflects other features like the two chromosome arms. To systematically account for this effect, we examined the first three eigenvectors for each chromosome in each replicate by correlating them with the gene density (compartmentalization is correlated with gene density, while other properties like chromosome arms generally are not). We required that PC1 show the highest correlation with gene density among the first three eigenvectors in every replicate. If this was not the case for a given chromosome, we eliminated that chromosome from all downstream analyses in all individuals to be conservative. Six chromosomes were eliminated in this way: chr1, chr9, chr14, chr19, chr21 and chr22. For the chromosomes that passed this filter, the sign of the first eigenvector (which is arbitrary) was adjusted such that the correlation between PC1 and gene density is positive, and this positive PC1 values correspond to compartment A. Finally, PC1 tracks were manually inspected to ensure that they are consistent with expected checkerboard patterns of compartmentalization.

#### 2.d. Directionality Index

Directionality Index was computed as previously described^16^. Briefly, upstream and downstream contacts within 2Mb window for each 40Kb bin were counted, and chi-square statistics were calculated under equal assumption. The sign of the chi-square statistics was adjusted such that positive values represent upstream biases. For some bins, there are more than five NA bins within 2Mb window and DI for those bins are not calculated. As noted in the main text, we made a slight variation of these DI scores for the QTL searches in which DI was recalculated using a window size of 200Kb to capture more local features.

#### 2.e. Insulation Score

Insulation scores were computed as previously described^16^with some adjustments. Briefly, contacts linking upstream and downstream 400Kb windows for each 40Kb bin were calculated in the O/E matrices instead of raw matrices. We further divided the contact frequency by the average of upstream and downstream 400Kb windows, to account for differences in contact density across the chromosome. The Insulation Scores were then ranged from 0 to 1, representing absolute insulation and no insulation respectively. Insulation scores for bins, for which more than 50% cells in the 400Kb window as NA values, were not computed. For the QTL search, we also calculated insulation scores using 200Kb window.

#### 2.f. TADs Calling

TADs were called using the same approach as described previously^16^. DI values for each 40Kb bins were used to build a Hidden Markov Model and predict the probability being upstream bias, no bias, and downstream bias. Regions switching from upstream bias to downstream bias were called as boundaries.

#### 2.g. FIRE

We first calculated FIRE score for each of 20 individuals, as described in our previous study^19^. Specifically, we mapped the raw reads to the reference genome hg19 as described above. Next, we removed all intra-chromosomal reads within 15Kb, and created 40Kb raw Hi-C contact matrix for each individual for each autosome. For each 40Kb bin, we calculated the total number of intra-chromosomal reads in the distance range of 15-200Kb. We then filtered bins as follows, starting from 72,036 autosomal 40Kb bins: First, we removed 40Kb bins with zero effective fragment size, zero GC content, or zero mappability score^64^. Next, we filtered out 40Kb bins within 200Kb of the bins removed in the previous step. We further filtered out 40Kb bins overlapping with the chr6 MHC region (chr6:28,477,797-33,448,354; hg19), which has extremely high SNP density that can make it difficult to correct for allelic mapping artifacts. This left 64,222 40Kb bins for downstream analysis. Next, we applied HiCNormCis^19^ to remove systematic biases from local genomic features, including effective fragment size, GC content and mappability. The normalized total number of cis intra-chromosomal reads is defined as FIRE score. We further performed quantile normalization across multiple individuals using R package “preprocessCore”. The final FIRE score is log transformation log2(FIRE score + 1) and converted into a Z-score to create a mean of 0 and standard deviation of one. To identify significant FIRE bins in each individual, we used one-sided P-value < 0.05. Ultimately, merging across all individuals, we identified 6,980 40Kb bins which are FIRE bin in at least one of 12 YRI individuals. Consistent with our previous findings^19^, we observed significant enrichment of GM12878 typical enhancers and super enhancers among these 6,980 40Kb FIRE bins (Supplemental Figure 1d). GREAT analysis^60^ further showed immune-related biological pathways and disease ontologies are enriched in these 6,980 40Kb FIRE bins (Supplemental Figure 1e).

### 3. Comparison of intra-individual vs inter-individual variation

To estimate variability between replicates, we computed Pearson correlation coefficient for all pairs of biological replicates for each score (DI, INS, FIRE and PC1). The pairs then can be divided into two groups based on whether they are from the same individuals as illustrated in Supplemental Figure 4c. We then tested if the distribution of Pearson correlation coefficients were different comparing two groups. Similar analysis was performed for contact matrices. For contact matrices, we calculated Pearson correlation coefficient for each distance and each chromosome separately as shown in Figure 1c.

### 4. Variable regions

#### 4.a. limma test for variable bins

To test regions that are variable across genomes, we applied limma^47^ with default parameters. First, values for each 40Kb bin in hg19 reference genome were calculated for each metrics tested (DI, FIRE, INS, PC1) as described above. DI, PC1, and INS scores were calculated based on contact matrices quantile normalized across 40 replicates. FIRE scores were calculated based on raw counts using HiCNormCis^19^ and then quantile normalized across 40 replicates. Second, we filtered out bins that are not testable. Specifically, FIRE scores were only tested for bins that are FIRE regions (p-value < 0.05) in any of 40 replicates. DI scores were only tested for bins where strong biases are observed (abs(DI) > 10.82757, which correspond to Chi-squared test p-value 0.001) in any of 40 replicates. INS scores were only tested for bins where strong insulation is observed (z-score transformed INS score < −1) in any of 40 replicates. No filterers were performed for PC1 scores. Third, we filtered out any bins that overlapping large SVs (> 10,000 bp) to avoid effect caused by SVs. Specifically, for FIRE, INS, and DI scores, bins that are within 200Kb, 400Kb, and 2Mb respectively upstream or downstream of large SVs were removed. For PC1 scores, bins overlapping large SVs were removed. Lastly, we applied limma standard model with individual as a fixed factor and eBayes correction. To estimate empirical false positive rate (FDR), we bootstrapped replicates to calculate the number of false positives in random background. Briefly, we random selected 40 or 22 replicates with replacement for LCL20 and YRI11 respectively, and identified variable regions as mentioned above. We performed 1,000 permutations and calculated empirical FDR as the average positive hits in 1,000 permutations divided by number of hits in real data.

#### 4.b. Normalizing Hi-C contact matrices using BNBC normalization

To directly compare individual Hi-C contact matrix cells across samples, we sought to remove unwanted per-cell variation owing to date of processing or other unknown ‘batch’ effects. To this end we developed Bandwise Normalization and Batch effect Correction (BNBC), described and evaluated in a separate manuscript (preprint on bioRxiv https://www.biorxiv.org/content/10.1101/214361v1). A brief description follows. For each chromosome and for each strata of distance between loci (a matrix “band”, hence the term “bandwise”), we correct for unwanted variation by taking the log counts-per-million-transformed values of all samples and generating a matrix whose entries are the observations for that chromosome’s matrix band across all samples (columns indexes samples and rows indexes contact matrix cells with anchor bins separated by a fixed distance). We then quantile normalize this matrix and regress out the impact of known batches (here, date of processing) using ComBat^65^ (specifically we correct both mean and variance). This procedure essentially conditions on genomic distance. We correct the majority of each contact matrix for each chromosome for each sample: we correct all but the 8 most distal matrix bands, for which we set all values to 0. The choice of the last 8 bands is empirical and reflects the small number of observations in each band matrix. The procedure is implemented in the bnbc package available through Bioconductor (http://www.bioconductor.org/packages/bnbc). Correction of contact matrices was performed on replicate-level data using the following LCLs: GM18486 (YRI-1), GM18505 (YRI-2), GM18507 (YRI-3), GM18508 (YRI-4), GM18516 (YRI-5), GM18522 (YRI-6), GM19099 (YRI-7), GM19141 (YRI-8), GM19204 (YRI-10), GM19238 (YRI-11), GM19239 (YRI-12), GM19240 (YRI-13), HG00731 (PUR-1), HG00732 (PUR-2), HG00512 (CHS-1), HG00513 (CHS-2). We note that NA19239 (YRI-12) replicate 1 and NA19240 (YRI-13) replicate 2 were excluded because the BNBC algorithm requires multiple samples from a given experimental batch to estimate batch effect parameters.

#### 4.c. Identifying biological variability in Hi-C contact matrices

To identify contacts with significant levels of between-individual variability we employed the following procedure, which mimics the analysis for INS, DI, FIRE and PC1, on contact matrices normalized by BNBC (see section 4b). For each contact matrix cell (representing loci separated by less than 28 Mb, this is a subset of the matrix cells normalized by BNBC) we used a linear model with individual modeled as a fixed factor, note we have 2 growth replicates for almost every individual. We used a parametric likelihood ratio test (equivalent to an F-test) to test whether there was significant between-individual variation. We used the IHW framework^49^ with the distance between anchor bins as informative covariate, to increase power and estimate false discovery rate. We used a FDR of 10% as significance threshold, resulting in 115,817 contact matrix cells with significant biological variability across the autosomes. To estimate effect size (depicted in Figures 2a, 3a and Supplementary Figure S5) we used a linear mixed effect model with individual as random effect, to decompose the variance into between-individual variability (biological) and within-individual variability (technical). As the measure of biological variability in these figures, we used the estimated biological variance. For this analysis, all 16 samples we normalized using BNBC were used.

#### 4.d. Correlation with other datasets

To examine correlation between 3D genome organization and other genome features, we reidentified variable regions with the same pipeline mentioned above using only individuals of which data is available for other gnome features, and then computed Spearman correlation coefficient between 3D genome metrics (DI, INS, PC1, and FIRE) and other genome features (RNA-seq, ChIP-seq, and DNase-seq) for each 40Kb bin that is variable. Signals for each 40Kb bins were calculated by averaging signals for the bin. Specifically, signals for ChIP-seq were the average signal of all peaks with in the bin, signals for RNA-seq were the average FPKM of all genes in the bin, and DNase signals were simply average signal for each base pair in the bin. In some cases, serval consecutive bins were identified as variable. We only kept the bin with strongest signal for other genome features among consecutive bins. To generate random backgrounds, we permutated individual labels for the same set of bins and recomputed Spearman correlation coefficient. 10,000 such permutations were used to calculate the statistical significance of departure from the null hypothesis in which the median value of true correlation values and permutated correlation values are equal. Similar analysis was performed for variable matrix cells with the following modifications. First, we used the variable matrix cells in the preceding section 4c. Second, to correlate matrix-cell-level contacts with bin-level DNase and ChIP-seq signals, anchor bins of variable matrix cells were used. Since each anchor bin may belong to more than one matrix cells, we only used each bin once and selected the one with the highest Spearman correlation coefficient. Exactly same approach was performed during permutation to ensure a fair comparison.

### 5. Phasing variants

Phasing of variants was performed based on HaploSeq pipeline^43^. Briefly, 1) Variants were filtered to keep only bi-allelic SNPs heterozygous in a given individual; 2) Aligned Hi-C bam files were realigned and recalibrated using GATK 3.4.0^66^ based on SNPs in the individual; 3) Filtered SNPs and realigned bam files were then used as input to run HAPCUT^67^; 4) Results from HAPCUT were further filtered to keep only the largest haplotype block and combined with homozygous alt SNPs as input for imputation using Beagle 4.0^68^ using 1000 Genome Phase 3 data excluding individual to phase as reference panel; 5) Results from Beagle were then combined with results of HAPCUT by removing conflicting phased SNPs. For all auto chromosomes except 1 and 9 in 18 out 20 individuals, we were able to obtain a single haplotype block. For chromosome 1 and 9, two arms were phased separately because of large heterochromatin region surrounding centromere. X chromosome was only phased for female individuals. We excluded NA19193 and HG00514 from phasing because of the lack of available high quality of genotypes. We evaluated accuracy of phasing in three probands in trios (NA19240, NA12878, HG00733) and found phasing results are of very high accuracy (~97.71%). Specifically, we calculated accuracy as percentage of correctly phased variants among total phased variants. Only variants whose transmission from parents can be ambiguously identified were used in calculation of accuracy where at least on parent is homozygous. Detailed statistics for phasing are listed in Supplemental Table 6.

### 6. CTCF motif variation and looping strength

GM12878 loops and motif positions were obtained from Rao *et al* 2014^14^ (GSE63525_GM12878_primary+replicate_HiCCUPS_looplist_with_motifs.txt.gz; N=9,448 HiCCUPS loops). We limited our analysis to autosomal *cis* loops in which a CTCF motif in one of the anchor regions overlaps a SNP (N=572). To evaluate the impact of motif disruption, we first identified eight “key” positions in the CTCF PWM (Jaspar MA0139.1)^69^ in which a single base has higher than 0.75 probability. We refer to SNPs at these positions in motif occurrences with one allele matching the high-probability base as “motif disrupting SNPs”. We refer to alleles matching the consensus base in the motif as strong motif alleles (S), and alleles matching any other base as weak motif alleles (W). There are N=142 loops with a motif disrupting SNP in a convergently-oriented CTCF motif, which refer to below as testable loops. For each testable loop, we extracted the Hi-C interaction frequency in the loop bin from each LCL, and classified as either “WW”, “SW”, or “SS” depending on the individual’s genotype at the corresponding motif disrupting SNP. To enable aggregation of data across different SNPs, we set the mean “SW” interaction frequency for each SNP to 1 and normalized all values for that SNP accordingly. These values are plotted in Figure 4b. In addition, for each testable loop we extracted a submatrix including the loop bin as well as 15 bins upstream and 15 bins downstream. Submatrices with missing values were discarded. For each SNP, we calculated the mean submatrix for each genotype, and then subtracted submatrices to calculate the difference in each matrix cell per “W” allele (i.e. SS-SW and SW-WW). These differences were then averaged across all SNPs and plotted in Figure 4c. Submatrices with missing values were discarded. For the allelic analysis in S/W heterozygous individuals, we used chromosome-span phasing results (see methods section 4) to split the Hi-C reads from each chromosome in each LCL into two separate haplotypes. Specifically, we required at least one base pair overlap with phased heterozygous SNPs with high base calling score (>13) and high mapping quality (>20). Reads overlapping indels or containing SNPs from both haplotypes were not used. Approximately 7.89% of Hi-C reads covered a heterozygous variant and could thus be assigned to one of the two haplotypes. The accuracy of haplotype assignment was evaluated by fraction of homologous-trans (h-trans) read, which contain SNPs from both haplotypes. On average ~1% reads were h-trans, suggesting high quality of the assignment. For each testable loop, we defined 40Kb windows around the center of each loop anchor region and calculated the number of reads connecting these two anchor windows (“loop reads”) on each haplotype. For each heterozygous LCL, we then calculated the percentage of loop reads that occur on the haplotype containing the S allele at the motif disrupting SNPs anchor. We required that at least 10 total loop reads were present for a given loop in a given heterozygous LCL, leading to a total of 218 data points from 105 different loops for inclusion in Figure 4d.

### 7. Identification of QTLs

#### 7.a. Testable bins

To identify testable bins for FIRE-QTL, DI-QTL, INS-QTL and C-QTL searches, we began with 72,036 autosomal 40Kb bins based on reference genome hg19. We eliminated “unreliable” bins with effective length, GC content, or mappability equal to zero^70^, resulting in 66,597 bins remaining. We further removed any 40Kb bins within 200Kb of an unreliable bin, resulting in 64,337 40Kb bins. We also removed bins covering the chr6 MHC locus (hg19: chr6:28,477,797-33,448,354, which is extremely polymorphic and may lead to complex mapping artifacts that are difficult to correct. To eliminate false signals in Hi-C data that could arise from large structural variations (SVs), we obtained SVs from the 1000 Genomes consortium^35^ (ftp://ftp-trace.ncbi.nih.gov/1000genomes/ftp/phase3/integrated_sv_map/ALL.wgs.integrated_sv_map_v2.20130502.svs.genotypes.vcf.gz) and removed bins which overlap one or more structural variants previously annotated in these individuals (N=123,015 SVs), or within 200Kb of large structural variations (>10Kb, N=1,253 SVs). These filtering steps yielded a set of 51,511 testable bins, which represent a common starting point for FIRE-QTL, DI-QTL, INS-QTL and C-QTL searches as described below.

#### 7.b. Testable SNPs

We began with a list of 15,765,667 variants among all 20 LCL individuals (Supplemental Table 3). We kept 14,177,284 variants among 11 unrelated YRI individuals, removed all indels, HindIII site polymorphisms, multi-allelic SNPs, and SNPs with minor allele frequency (MAF) < 5%. We also required that remaining SNPs were within the 51,511 testable bins described above, and that both alleles were present in at least 2 individuals in the discovery set individuals. (N=4,132,791 SNPs remaining). Finally, where multiple SNPs in the same bin were in perfect LD among 11 unrelated YRI individuals, we selected one with the smallest genomic position (to avoid the introduction of a random selection that would not be perfectly reproducible), ultimately yielding 1,304,404 potentially testable SNPs that served as a common input set to all QTL searches.

#### 7.c. Power Calculations

To explore the power of our approach and data, we performed a Monte Carlo-based power calculation. Specifically, we varied four variables: (1) the minor allele frequency of a variant; (2) the effect size of genotype (a fixed effect); (3) the variability between subjects (a random effect); (4) the variability of the residuals. For contact QTLs, we also varied the mean of the Hi-C contact frequency in question. For analyses reported, we fixed the number of replicates-per-subject to be 2 (consistent with our study design). We explored a variety of settings for these parameters to assess power as each variable changes (see Supplemental Table 7). Each setting tested was chosen to reflect the distribution of observed values in our real Hi-C data. For each configuration of parameters, we performed the following simulation: We simulated genotypes by randomly sampling a set of alleles (one allele per subject) from a binomial distribution parameterized by the number of subjects and the MAF; we repeated this process twice and create per-subject genotypes by adding the results of the sampling of alleles. We simulated per-subject random effects, and per-sample residuals. To obtain a given sample’s simulated Hi-C contact matrix value, we added the mean Hi-C contact matrix value to that sample’s simulated genotype (multiplied by the pre-specified effect size), the specific subject’s random intercept and the sample’s random residual. After performing this for all samples, we then fitted the same LMM model used in our QTL search. We repeated this simulation and model fitting process 1,000 times and computed power as the fraction of times the null hypothesis that the effect of genotype is equal to 0 is rejected at a nominal p-value of 0.05.

#### 7.d. FIRE, DI, and INS QTL searches. 7.d.i. FIRE tested bins and SNPs

We limited our FIRE QTL search to the subset of testable bins that were called as FIRE in at least one YRI LCL (N=5,822 FIRE test bins), and the subset of testable SNPs therein (N=128,137 FIRE test SNPs). **7.d.ii. INS tested bins and SNPs**. For the INS-QTL search, we examined 328,530 test SNPs with 12,976 variable INS bins (see methods section 4a). **7.d.iii. DI tested bins and SNPs**. For the DI-QTL search, we examined 181,950 test SNPs with 7,590 variable DI bins ((see methods section 4a). For the DI-QTL search, we further classified each DI bin based on which whether it showed stronger upstream or downstream bias, because we saw a Simpson’s paradox when we considered them together (see discussion above, Supplemental Figure 10b). This was done as follows: for each bin we evaluated the DI score in each of 11 unrelated YRIs and identified the DI score among these individuals with the largest absolute value. We defined a bin as “upstream DI bias” if the DI score with the highest absolute value was positive, or “downstream DI bias”, if the DI score with the highest absolute value was negative. Only 37/7,590 bins (0.4%) had individuals with both positive and negative DI. **7.d.iv. LMM QTL searches**. For each test SNP, we identified the 40Kb bin it belongs to, and fitted a linear mixed effect model, using FIRE, DI (200Kb window; see section 2d), or INS score (200Kb window; see section 2e) in each biological replicate as the response variable and genotype of that testable SNP as the explanatory variable. Since two biological replicates from the same individuals are correlated included an individual-specific random effect to account for within-individual correlation. We used the R package “nlme” and R function “gls” to fit the linear mixed effect model. The quantile-quantile plots (QQplot) showed only minor genomic inflation (median p-value = 0.4821, lambda = 1.0864 for FIRE-QTLs; median p-value = 0.4864, lambda = 1.0649 for upstream-biased DI-QTLs; median p-value = 0.4828, lambda = 1.0826 for downstream-biased DI-QTLs; median p-value = 0.4865, lambda = 1.0646 for INS-QTLs). The linear mixed effect model identified 476, 315, 315, and 1,092 SNPs with false discovery rate (FDR) less than 0.20 for FIRE, upstream-biased DI, downstream-biased DI and INS, respectively. When more than one SNP in the same bin was identified, we selected the SNP with lowest p-value among them to be included in the final QTL sets. After this filtering, we ended up with 387 candidate FIRE-QTLs, 268 candidate upstream-biased DI-QTLs, 277 downstream-biased DI-QTLs, and 911 candidate INS-QTLs. As a control for each of these QTL searches, we randomly shuffled the score in question (i.e. FIRE, DI, or INS) among all 11 YRI individuals and performed QTL searches on this permuted data. In each of these tests, we found no SNPs associated with the permuted scores at FDR < 0.20.

#### 7.e. C-QTL search

To find QTLs affecting Hi-C contact strength we first identified 115,187 Hi-C contact matrix cells exhibiting substantial biological variability as described in section 4b, and constrained our QTL search to these cells. We then intersected these contact cells with 1,304,404 testable SNPs by requiring a SNP to sit in one anchor bin of one of these variable matrix cells. We also filtered out matrix cells to ensure both anchor bins of the matric cell are among 51,511 testable bins. In total, we obtained 3,109,039 tests involving 687,655 SNPs and 54,880 matrix cells on all 22 autosomes. For each test, we used the BNBC normalized data described in section 4, but used only the 11 unrelated YRI individuals with genotypes available and fit a linear mixed effect model in which genotype is a fixed effect and subject is a random intercept. We then used “lmerTest” package in R to estimate p-values for the fixed effect of genotype^71^. We used the IHW framework^49^ to estimate FDR, with the distance between anchor bins as an informative covariate, and call any matrix cell with FDR < 0.2 as significant. We further filtered significant tests by selecting the most significant SNPs per matrix cell and kept the leftmost SNPs among SNPs in perfect LD in two anchor bins of the matrix cell. After filtering, we ended up with 463 tests involving 345 SNPs and 463 matrix cells. To make the aggregate contact plots in Figure 4g, we recoded the genotypes based on the direction of effect such that 0,1, 2 refer to the genotypes containing 0, 1 or 2 alleles associated with the increased phenotype, respectively. Next, to avoid aggregating the same submatrix multiple times, we filtered by 1) selecting only the most significant matrix cell associated with each QTL, 2) selecting only the most significant QTL associated with each anchor bin (in some cases the same bin anchors multiple matrix cells associated with different QTL SNPs). This filtering left 165 unique matrix cell QTL interactions for plotting. For each matrix cell, we then extracted a submatrix including 25 bins upstream and 25 bins downstream. Submatrices with missing values were discarded. For each QTL, we then calculated the mean submatrix values for each genotype, and then subtracted submatrices to calculate the difference in interaction frequency between the 1 and 0 genotypes, and between the 1 and 2 genotypes. These differences were then averaged across QTLs and plotted in Figure 4g

#### 7.f. Validation of QTLs in additional individuals

Our validation set included six unrelated individuals not included in the discovery set: NA12878, NA19240, HG00512, HG00513, HG00731 and HG00732. For each QTL, we collected the genotype among six additional individuals, and the corresponding FIRE, DI, or INS scores. Note that a small fraction of QTLs have missing genotypes in these six individuals (coded as “-1”), and these missing data points were eliminated from validation analysis. We examined the distributions of scores for each genotype. For each QTL type (i.e. FIRE, DI, or INS), we found that the same direction of effect observed in the discovery set is observed on average in the validation set. To assess the significance of this observation, we approximated the null expectation as follows. For FIRE-QTLs, for example, we started from all 128,137 FIRE test SNPs and 5,822 FIRE test bins. Note that in our discovery set, we identified 387 FIRE-QTLs, each in a different 40Kb bin. To create a random control SNP group, we first randomly selected 387 40Kb bins from all 5,822 FIRE test bins. Next, within each select bin, we randomly selected one SNP, and combined all these 387 selected SNPs into a control SNP group. We then tested their SNP effect on the six additional individuals. We repeated such sampling with replacement 1,000 times, to create a null distribution of positive and negative SNP effect, respectively. We performed the same type of permutations for DI, INS. Similar analysis was performed for C-QTLs with a few modifications. First, we only used replicates from NA19420, HG00512, HG00513, HG00731 and HG00732 as explained in methods section 4. Second, 1,000 random permutations were performed by sampling matrix cells instead of bins. Third, we used values of biological replicates separately instead of as merged data because the BNBC normalization is performed at the level of replicates.

#### 7.g. Examining epigenetic variation at FIRE, DI, INS, and C-QTLs

To examine epigenetic variation at 3D genome QTLs, we re-analyzed DNase-seq data from 59 LCLs^38^, histone modification ChIP-Seq data (H3K27ac, H3K4me1 and H3K4me3) for 65 LCLs^39^, and CTCF ChIP-seq data from 11 LCLs^54^. These data were re-mapped using the WASP pipeline to control for allelic mapping artifacts and calculating the signal in 40kb bins as described above in section 3.b. We examined the effect of genotype at FIRE, DI, INS or C-QTLs on DNase-seq and ChIP-seq signal by linear regression. As a control, we randomly selected the matched number of SNPs with the same approach described in section 7.f and re-did such validation analysis. We repeated such random sample 1,000 times to create the empirical null distribution of no genetic effect. For C-QTLs, we used the sum of epigenetic features in two anchor bins to calculate correlation with contact frequency.

### 8. Nominal fraction analyses

#### 8.a. Comparing between 3D chromatin QTL types

To compare between different 3D chromatin QTLs, we took the raw test results for each QTL set and projected other 3D QTLs into the test results. For example, in Figure 4j we selected subset of SNPs that are DI-QTLs and plotted them (dark green dots) using p-values from FIRE-QTLs along with all tested in the FIRE-QTL search (black dots). We also used all tested SNPs in the DI-QTL search (light green dots) as a control set. To assign significance to the overlap, we compared the fraction of SNPs with nominal significance (p-value<0.05) in each set: 1) DI-QTL tested SNPs that were not significant QTLs, and 2) DI-QTLs. We calculated p-values for this comparison by Chi-square test. To rule out the effect of sampling bias when selecting a small number of SNPs, we also performed permutation. In each permutation, we randomly selected the same number of SNPs as the real QTL set (from the full set of tested SNPs) and calculated the fraction with nominal significance. We then computed bootstrap p-values using 10,000 such permutations under the null hypothesis that the fraction of nominal significance is the same between QTLs and random selected SNPs. For C-QTLs, one SNP may be tested against multiple matrix cells, so we only keep the most significant p-value for each SNP to avoid biases towards SNPs with multiple tests.

#### 8.b. Comparing 3D chromatin QTLs to other molQTLs

Similar approaches were used to assess overlap between 3D chromatin QTLs and other molQTLs. We obtained full test results (all tested SNPs with the p-values) from previous molQTL studies and projected 3D chromatin QTLs into those test results. We the calculated fraction of nominal significance and used chi-square test to evaluate significance between 3D-QTLs and non-3D-QTLs. Similarly, we performed bootstrap to estimate significance empirically. One modification is that we extended our QTL sets by incorporating all SNPs in perfect LD with the same 40Kb bin because we may not use the same tagging SNP in our study as used in other studies. To ensure a fair comparison, we performed the same extension for the control sets of all tested SNPs.

#### 8.c. Comparing 3D chromatin QTLs to GWAS

Comparison with the GWAS results was performed in the same manner as described above in 8.b. for other molQTLs. Instead of test results for other molQTLs, we used summary statistics from previous GWAS.

### 9. FISH

#### 9.a. Cell preparation for FISH

Approximately 100,000 cells were adhered to center of PDL-coated coverslips (Neuvitro, GG-22-15-PDL) by placing 100 uL of cells at 1 x 10^6^ cells/mL. Cells on coverslips were incubated for an hour at 37°C, carefully washed with PBS, and fixed with 4% paraformaldehyde in 1X PBS for 10 mins. PFA was quenched with 0.1 M Tris-Cl, pH 7.4 for 10 mins, washed with PBS, and stored in 1X PBS at 4°C for up to 1 month.

#### 9.b. BAC probe labeling and preparation

All BAC clones were ordered from the BACPAC Resource Center at the Children’s Hospital Oakland Research Institute: “U” probe is RP11-74P5, “C” probe is RP11-337N12, and “D” probe is RP11-248M23. BAC DNAs were labeled with either Chromatide Alexa Fluor 488-5 dUTP (Invitrogen, C-11397) or Alexa Fluor 647-aha-dUTP (Invitrogen, A32764) using nick-translation kit (Roche, 10976776001), and incubated in 15°C for 4 hours. The nick-translation reaction was deactivated using 1 uL of 0.5 M EDTA, pH 8.0 and heated for 10 mins at 65°C. The probes were then purified using illustra ProbeQuant G-50 Micro Columns (GE Healthcare, 28903408) and eluted to a concentration of 20 ng/uL. Probes were mixed with Human Cot-1 DNA (Invitrogen, 15279011) and salmon sperm (Invitrogen, 15632011), and precipitated with 1/10th volume of 3M sodium acetate, pH 5.2 and 2.5 volume of absolute ethanol for at least 2 hours at −20°C. Probes were then spun down, washed with cold 70% ethanol, resuspended in formamide and 40% dextran sulfate in 8X SSC, and incubated at 55°C.

#### 9.c. Hybridization

Cells on coverslips were blocked with 5% BSA and 0.1% triton-X 100 in PBS for 30 mins at 37°C, and washed twice with 0.1% triton-X 100 in PBS for 10 mins each with gentle agitation at room temperature. Cells were permeabilized with 0.1% saponin and 0.1% triton-X 100 in PBS for 10 mins at room temperature. Next, they were incubated in 20% glycerol in PBS for 20 mins, freeze-thawed three times with liquid nitrogen, and incubated in 0.1M hydrogen chloride at room temperature for 30 mins. Cells were further blocked for 1 hour at 37°C in 3% BSA and 100 ug/mL RNase A in PBS. Cells were permeabilized again with 0.5% saponin and 0.5% triton-X 100 in PBS for 30 mins at room temperature. Lastly, they were rinsed with 1X PBS and washed with 2X SSC for 5 mins. For hybridization of probes, the prepared probes were denatured at 73°C for 5 mins in water bath. Cells were denatured in a two-step process in a 73°C water bath: 2.5 mins in 70% formamide in 2X SSC and 1 min in 50% formamide in 2X SSC. Denatured probes were transferred onto microscope slides, and coverslips were placed on top with cell-side facing down. The coverslips were sealed with rubber cement and incubated overnight at 37°C in a dark, humid chamber. Next day, coverslips were carefully removed and transferred onto a 6-well plate. Cells were washed at 37°C with gentle agitation, twice with 50% formamide in 2X SSC for 15 mins and three times with 2X SSC for 5 mins. The cells were then stained with DAPI (Invitrogen, D1306), mounted on microscope slides with ProLong Gold Antifade Mountant (Invitrogen, P36930), sealed with nail polish, and imaged.

#### 9.d. Microscope and analysis

Images were acquired with DeltaVision RT Deconvolution Microscope at UC San Diego’s department of neuroscience (acquired with award NS047101). Captured images were processed using the TANGO^72^ plugin in ImageJ for quantitative analysis. Each FISH experiment contained two probes labeled with different color dyes (either U-C or C-D). We limited our analysis to nuclei containing 2 labeled foci for each color (4 total foci), allowing us to more confidently distinguish foci *in cis* from those *in trans*. Distances were measured from the center of one color focus to the center of the closest focus of the other color.

### 10. Re-analysis of public datasets

#### 10.a. Analysis of ChIP-seq data from Kasowski et al and McVicker et al

Raw fastq files were downloaded from SRA database for each experiment (SRP030041 and SRP026077, respectively). Reads were aligned to hg19 reference genome using BWA MEM (Kasowski) or BWA ALN^63^ v0.7.8 (McVicker) with WASP pipeline^45^ to eliminate allelic mapping bias. Only reads with high mapping quality (>10) were kept. PCR duplicates were removed using Picard tools v1.131 (http://broadinstitute.github.io/picard). MACS2^73^ v2.2.1 was then used to call peaks using corresponding input files. For CTCF and SA1, default parameters were used for MACS2. For H3K27ac, H3K4me1, and H3K4me3, peak calling was done using “--nomodel” parameter because we do not expect sharp peaks for histone modifications. For H3K27me3 and H3K36me3, peak calling was done using “--nomodel --broad” parameter. Bigwig files were generated by MACS2 using fold enrichment for viewing in genome browser. All Kasowski data were processed in pair-end mode and both replicates were merged for analysis. All McVicker data were processed in single-end mode, and the pooled input data were used for all samples because there are no individual input files. To compute signals in peaks, we used a set of merged peaks across all individuals for each mark.

#### 10.b. Analysis of RNA-seq data from Kasowski et al

Raw fastq files were downloaded from SRA database (SRP030041). Reads were aligned to hg19 reference genome using STAR^74^ v2.4.2a with the WASP pipeline in pair-end mode to eliminate allelic mapping bias. Gencode^75^ v24 annotation was used to construct STAR index and computing FPKM. Only uniquely mapped reads were kept. Cufflinks^76^ v2.2.1 was applied to compute FPKM values. Both replicates were merged for analysis.

#### 10.c. Analysis of DNase-seq data from Degner et al

Raw fastq files were downloaded from SRA database for each experiment (SRP007821). Reads were aligned to hg19 reference genome using BWA ALN with the WASP pipeline in single-end mode to eliminate allelic mapping bias. Only reads with high mapping quality (>10) were kept. PCR duplicates were removed using Picard tools. Bigwig files were generated using makeUCSCfile commands in homer tools^77^ v4.9.1.

#### 10.d. Analysis of ChIP-seq data from Ding et al

Raw fastq files were downloaded from SRA database for each experiment (SRP004714). Reads were aligned to hg19 reference genome using BWA MEM v0.7.8 with the WASP pipeline to eliminate allelic mapping bias. Only reads with high mapping quality (>10) were kept. PCR duplicates were removed using Picard tools. We performed quality control for CTCF ChIP-seq data by FRIP (Fraction of Reads In Peaks) and used datasets with FRIP > 10. Bigwig files were generated using bamCoverage commands in deepTools^78^ v2.3.3. To compute signals in peaks, we used the merged CTCF peaks from Kasowski data.

#### 10.e. Analysis of ChIP-seq data from Grubert et al

Bigwig files and peaks for H3K27ac, H3K4me1 and H3K4me3 were downloaded from GEO database (GSE62742). Peaks for each mark were merged and then used to compute the averaged signal.

## DATA AVAILABILITY

All raw sequencing data and many processed data files are available through NCBI’s Gene Expression Omnibus (GEO) (GSE128678), as well as through the 4D Nucleome data portal (https://data.4dnucleome.org). Additional processed data not provided above in supplement, as well as code, are available on the Ren lab’s website at http://renlab.sdsc.edu/renlab_website//download/iqtl/, or by request.

## ACKNOWLEDGEMENTS

This work was supported by NIH grant U54DK107977 to B.R. and M.H. D.U.G. was supported by fellowships from the A.P. Giannini Foundation and the NIH Institutional Research and Academic Career Development Awards (IRACDA) program. The authors would like to acknowledge members of the Ren lab, and Dr. Graham McVicker for important discussions and feedback during preparation of this manuscript. B.R. is co-founder and share holder of Arima Genomics. A.D.S. is employee and share holder of Arima Genomics.

## SUPPLEMENTAL FIGURE LEGENDS

**Supplemental Figure 1. Hi-C derived molecular phenotypes measured across 20 LCLs.** (a) Hi-C contact matrices show for all 20 LCLs. For comparison, we also show data from H1 human embryonic stem cells (H1-ES), and 4 lineages derived from H1 by in vitro differentiation^21^ (H1-ME = Mesendodermal cells, H1-NPC = Neuronal precursor cells, H1-TR = Trophoblast-like cell, MSC = Mesenchymal stem cells). These H1-derived cell line represent different cell types from the same individual (i.e. same genetic background). The region shown here is the same as Figure 1a (chr8:125,040,000-132,560,000; hg19). (b) Same region as above, but showing PC1 and FIRE values. ChIP-seq data for several histone modifications, CTCF, and Cohesin subunit SA1 are shown for one LCL (YRI-13, GM19240) as a reference for the epigenomic landscape^42^. (c) Same region as above, but showing DI and INS values. (d) Bar plots show the percentage of super-enhancers (left) or typical enhancers (right) in GM12878^59^ that overlap with 6,980 LCL FIRE bins (called as FIRE in at least one individual in our dataset) and 6,980 random 40kb bins. (e) Biological Process Gene Ontology terms associated with genes proximate to FIRE regions as defined by GREAT^60^.

**Supplemental Figure 2. FIRE measures density of local interactions.** Illustrative example showing that overall density of Hi-C reads (all reads irrespective of location of interacting partner, all *cis* interactions, or all *trans* interactions) is highly consistent across the genome. However, interactions between partners separated by 15-200kb (“FIRE” distance) show enrichment in regions of the genome with marks of regulatory and transcription activity (H3K4me1, H3K4me3, H3K36me3 from ENCODE for CEU-1 / GM12878 shown for reference)^61^. ChromHMM functional annotations for CEU-1 / GM12878 are also shown^62^. pr-a = active promoter, pr-w = weak promoter, pr-i/p = inactive/poised promoter, en-s = strong enhancer, en-w/p = weak/poised, ins = insulator, tr-el/tr = transcriptional elongation or transcriptional transition, tr-w = weak transcribed, rep-p = polycomb-repressed, het/rep/cnv = heterochromatin, low signal, repetitive, or copy number variation. Top panel (a) shows the long arm of chr14 (chr14:24,406,737-104,693,368; hg19). Bottom panel (b) is a zoomed-in view of region boxed by dotted lines above (chr14:58,000,000-63,500,000; hg19).

**Supplemental Figure 3. Aggregate looping interactions in each sample.** Aggregate plots show the interaction frequencies at GM12878 HiCCUPS loops from Rao et al 2014 in each sample examined here. All autosomal *cis* loops with anchor bins separated by more than 40kb are included here (N=8,893). The middle bin represents the interaction frequency between two 40kb bins containing loop anchor bins. The full submatrix extends 10 bins (400kb) upstream and downstream of the interaction bin (x and y axis). The color scale indicates the average interaction frequency per loop from Hi-C contact matrices.

**Supplemental Figure 4. 3D chromatin variation among 20 LCLs and H1-derived lineages.** (a) Graphical representation of the shuffling scheme used to assess biological variability in Figure 1b-d, and here in panels b-e. (b)-(e) Boxplots show Pearson correlation coefficient between biological replicates from the same cell line (Replicates = “True”), and between replicates from difference cell lines (Replicates = “Shuff”; short for “shuffled”). The set of cell lines considered is indicated below each box. LCL = 20 LCL cell lines (40 replicates); H1 = H1-ES and the four derived lineages H1-ME, H1-NPC, H1-TB, H1-MSC (5 cell lines, 10 replicates); H1+LCL = 20 LCLs and 5 H1-derived lines considered together (50 replicates). All phenotypes examined here (DI, PC1, INS, or FIRE) show a signature of cell-type specificity whereby they are more similar across individuals when looking at the same (or highly similar) cell types (i.e. LCLs), relative to comparing across cell types within an individual (i.e. same genetic background, H1s). Statistical significance calculated by two-sided Wilcoxon rank sum test. (f) Dendrograms from hierarchical clustering of 40 Hi-C replicates based on one of four Hi-C-derived phenotypes, as indicated above each dendrogram (DI, PC1, INS, or FIRE). In the most of cases, replicates from the same cell lines cluster together. (g) Principal Component Analysis of 20 LCLs using one of four Hi-C-derived phenotypes, as indicated above each plot. Each population in our study is represented by a different color as indicated in the color key to the right. Note that the LCLs tend to separate by population in each plot.

**Supplemental Figure 5. Characterization of variable regions of 3D chromatin conformation.** (a) Same region as in Figure 2a (chr15:94,280,000-99,280,000), but showing reproducible variation in PC1, and full square matrices for contact matrix variability as opposed to the half-matrices shown in 2a. FISH probes used in Figure 2 are represented as grey boxes to the top and right of square heatmaps. (b) Similar to Figure 2b, but with additional data columns. The “merged regions” column shows the number of regions after merging variable bins that are immediately adjacent to each other. The empirical false positives and FDR columns show the number and percentage of false positive variable regions detected in 1000 permutations with shuffled labels. (c) Venn diagrams showing the overlap of variable regions identified using either all 20 LCLs (“LCL20”) or only the 11 unrelated YRI LCLs (“YR111”). Only bins tested for both LCL20 and YRI11 are included here. (d) Venn diagrams showing the number of variable bins for each phenotype or combination of phenotypes. The diagram on the left includes bins that were testable for at least one of the four phenotypes. The diagram on the right only includes bins that were testable for all four phenotypes. (e) Mosaic plots show the significance of overlaps between variable regions in a pairwise fashion. P-values calculated by two-sided chi square test.

**Supplemental Figure 6. Additional characterization of variable regions of 3D chromatin conformation.** (a) Same underlying FISH data as in Figure 2e, but here comparing the distance between U and C probes to the distance between C and D probes within the same LCL. P-values calculated by two-sided Wilcoxon rank sum test. (b) As in Figure 2d, blue line shows the fraction of variable matrix cells distributed across a range of interaction distances. Black line shows the fraction of all matrix cells distributed across the same range of interaction distances. (c) Mosaic plots show the significance of overlap between variable regions and anchor bins of variable matrix cells. P-values calculated by two-sided chi square test.

**Supplemental Figure 7. Coordinated variation between 3D chromatin conformation and multiple molecular phenotypes.** (a) Same region as in Figure 3a (chr6:126,280,000-131,280,000; hg19), but showing additional individuals and additional data types as indicated. (b) Representation of permutation scheme used to calculate P-values in panels c as well as in Figure 3b and Supplemental Figures 8 and 9. (c) Density plots in the top left quadrant show Spearman correlation coefficients (SCC) between PC1 and molecular phenotypes as indicated in the top margin of panel. Density plots in the top right quadrant show the same underlying data but plotted as absolute SCC to highlight the shift of real correlations (red lines) towards one, relative to the permutated values (grey). We show both SCC and abs(SCC) because SCC is useful to observe bias toward positive or negative correlations, but the bootstrap p-values are calculated based on the abs(SCC) to reflect correlations that are more extreme (positive or negative) than expected based on approximation of the null hypothesis. For the top left and top right quadrants, the ChIP-seq and RNA-seq data in the figure from Kasowski *et al*, 2013^42^ (same as in Figure 3b), with the addition of histone modifications H3K4me1, H3K4me3, and H3K36me3, as well as DNase-seq data from Degner *et al*, 2012^38^. ChIP-seq, RNA-seq, and DNase-seq plots include eight, seven, and eleven individuals, respectively (i.e. all individuals for which both Hi-C and the data type in question are available). Bottom left quadrant shows SCC like above, but using variable regions called in only 11 individuals. Bottom right quadrant shows SCC using ChIP-seq data from McVicker *et al*, 2012^40^. These plots include ten individuals for which ChIP-seq and Hi-C data are available. In all cases, p-values are calculated as the number of permutations with a mean absolute SCC greater than the observed values.

**Supplemental Figure 8. Correlations between DI, INS and multiple molecular phenotypes.** Similar schema to Figure 7c, but focusing on DI in (a), and INS in (b).

**Supplemental Figure 9. Correlations between FIRE, interaction frequency and multiple molecular phenotypes.** Similar schema to Figure 7c, but focusing on FIRE in (a), and contact frequency (examining the anchor bins of variable matrix cells) in (b).

**Supplemental Figure 10. 3D chromatin QTLs.** (a) QQ plots for each QTL search. In each QQ plot, the X-axis is the −log10 theoretical quantiles calculated from the uniform distribution. The Y-axis is the −log10 p-value calculated from linear mixed effects model for each type of QTL search. The grey area represents the 5% - 95% confidence bands based on Beta distribution Beta(i, M-i+1), where i is the i th order statistics and M is the total number of tested SNPs. (b) Genotype trend for bins with positive DI (left), negative DI (right), and all QTLs (right). A Simpson’s paradox is observed when all bins are considered together. (c) Number of direct overlaps between QTL sets. (d) Similar schema to Figure 4e, but showing FIRE, INS, DI score (indicated on the Y axis) as a function of genotype for each QTL set as indicated above each column. Grey boxes highlight the cases plotted in Figure 4e where the signal type and QTL set are the same.

**Supplemental Figure 11. Influence of 3D chromatin QTLs on epigenomic and disease phenotypes.** (a) Similar schema to Figure 5a, but showing DI-QTLs in positive bins (top) and negative DI bins (bottom). (b) Left subpanel shows the enrichment for 3D QTL SNPs with nominal significance in the indicated epigenetic or eQTL study calculated as follows: (fraction of indicated 3D QTL SNPs with nominal significance in the indicated molQTL study) / (fraction of SNPs tested in the indicated 3D QTL search with nominal significance in the indicated molQTL study). Asterisks mark values with *p*<0.05 by chi-square test (middle panel), and permutation test (right panel). Right panel shows the proportion of 1,000 random subsets selected from the tested SNPs with enrichment values higher than the indicated true QTL set. Dotted lines mark *p*=0.05. (c) QQ plot shows the results of H3K4me1 QTL search from Grubert et al., with all tested SNPs shown as black points, and two subsets as follows: SNPs also tested in our C-QTL search (light blue), and SNP called as C-QTLs or in perfect LD with C-QTLs in the same 40Kb bin (dark blue).

## TABLES

Supplemental Table 1. LCLs included in the study.

Supplemental Table 2. Public datasets re-analyzed in this study.

Supplemental Table 3. Hi-C mapping statistics.

Supplemental Table 4. Regions showing evidence of biological variability in 3D chromatin conformation.

Supplemental Table 5. Matrix cells showing evidence of biological variability.

Supplemental Table 6. Summary of phasing results.

Supplemental Table 7. Power calculations.

Supplemental Table 8. 3D chromatin conformation QTLs.

Supplemental Table 9. Overlaps between 3D chromatin QTL and GWAS catalog.

