## Supplemental Figures 1-11 for "Common DNA sequence variation influences 3-dimensional conformation of the human genome"

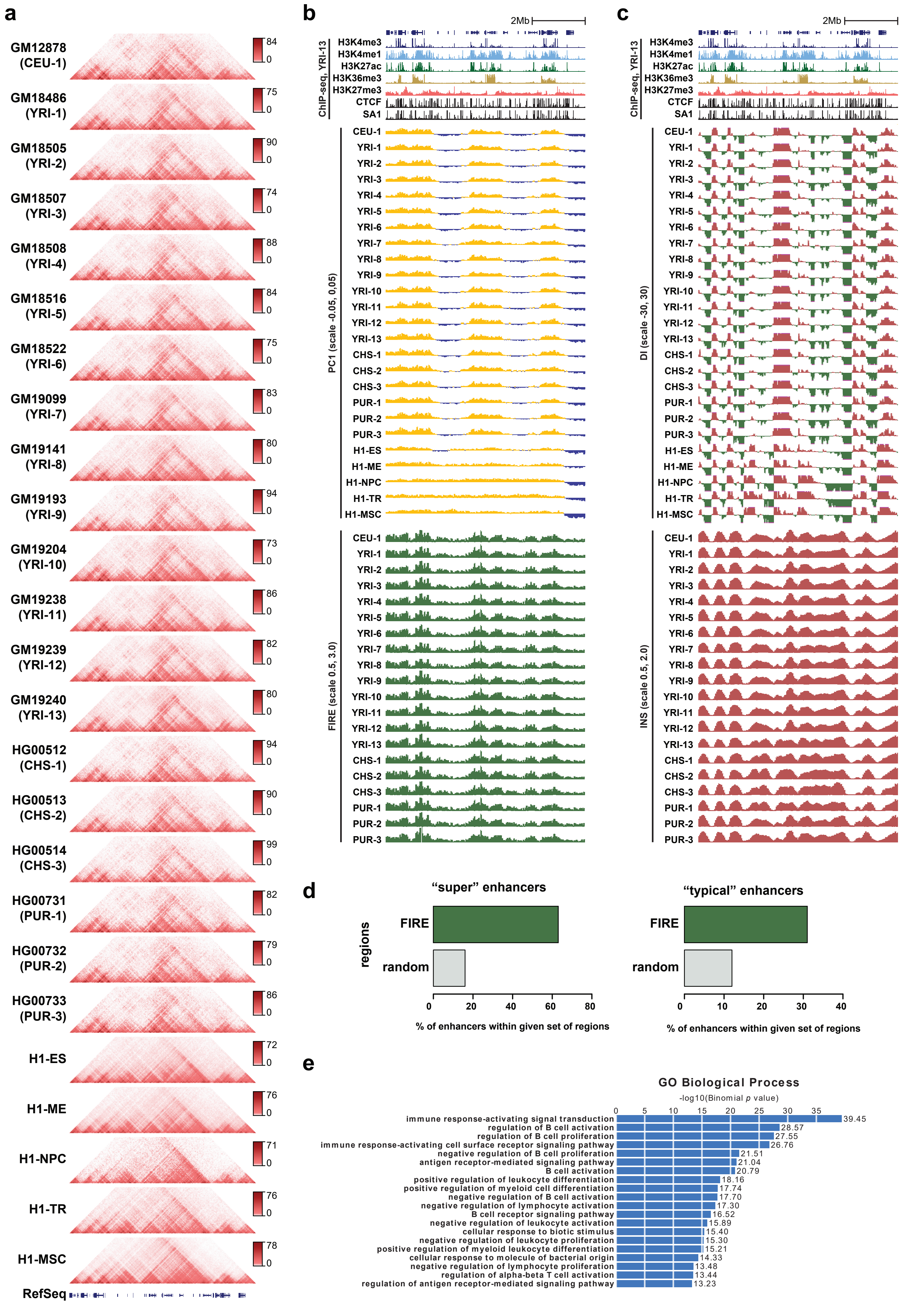


Supplemental Figure 1

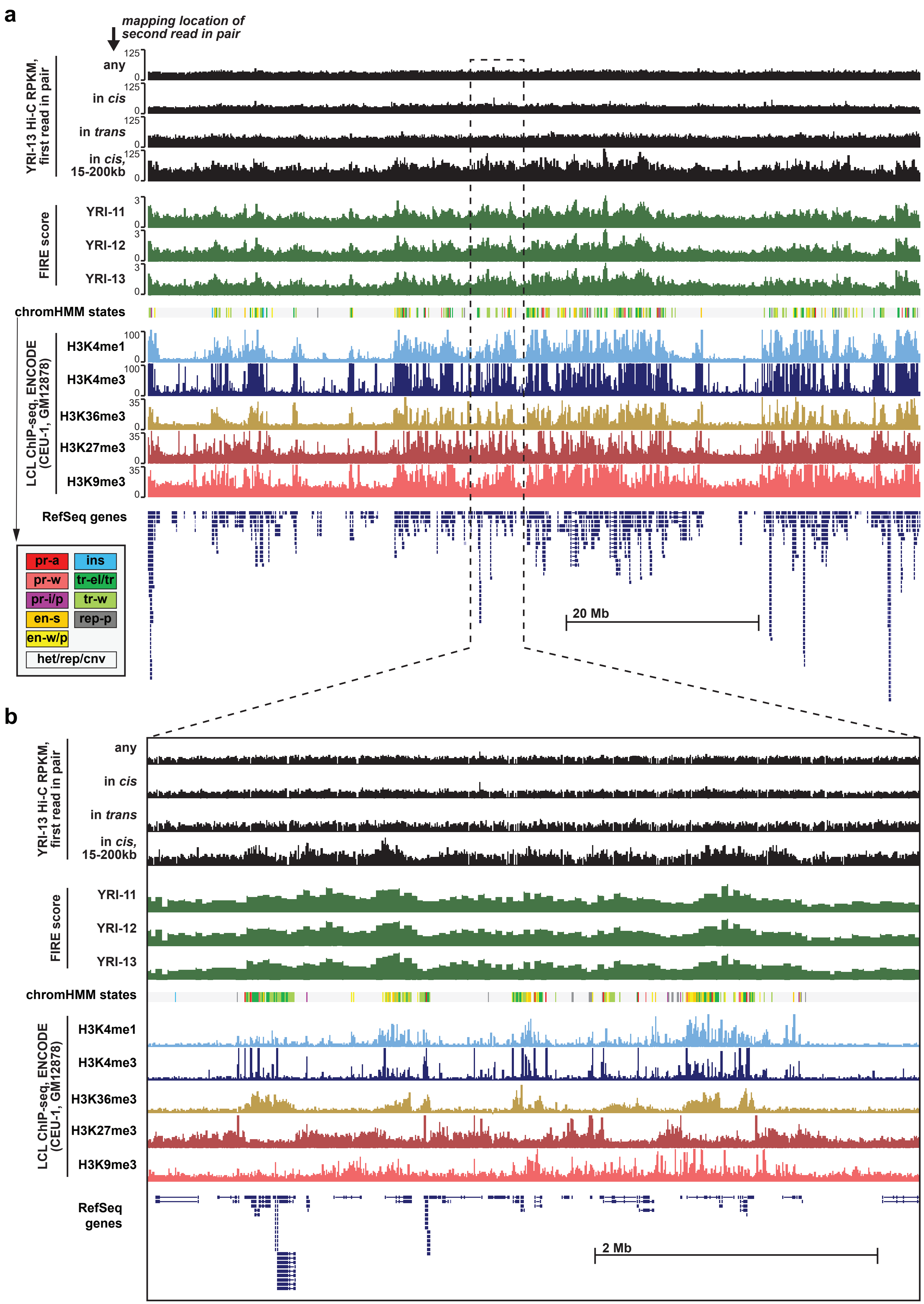


Supplemental Figure 2

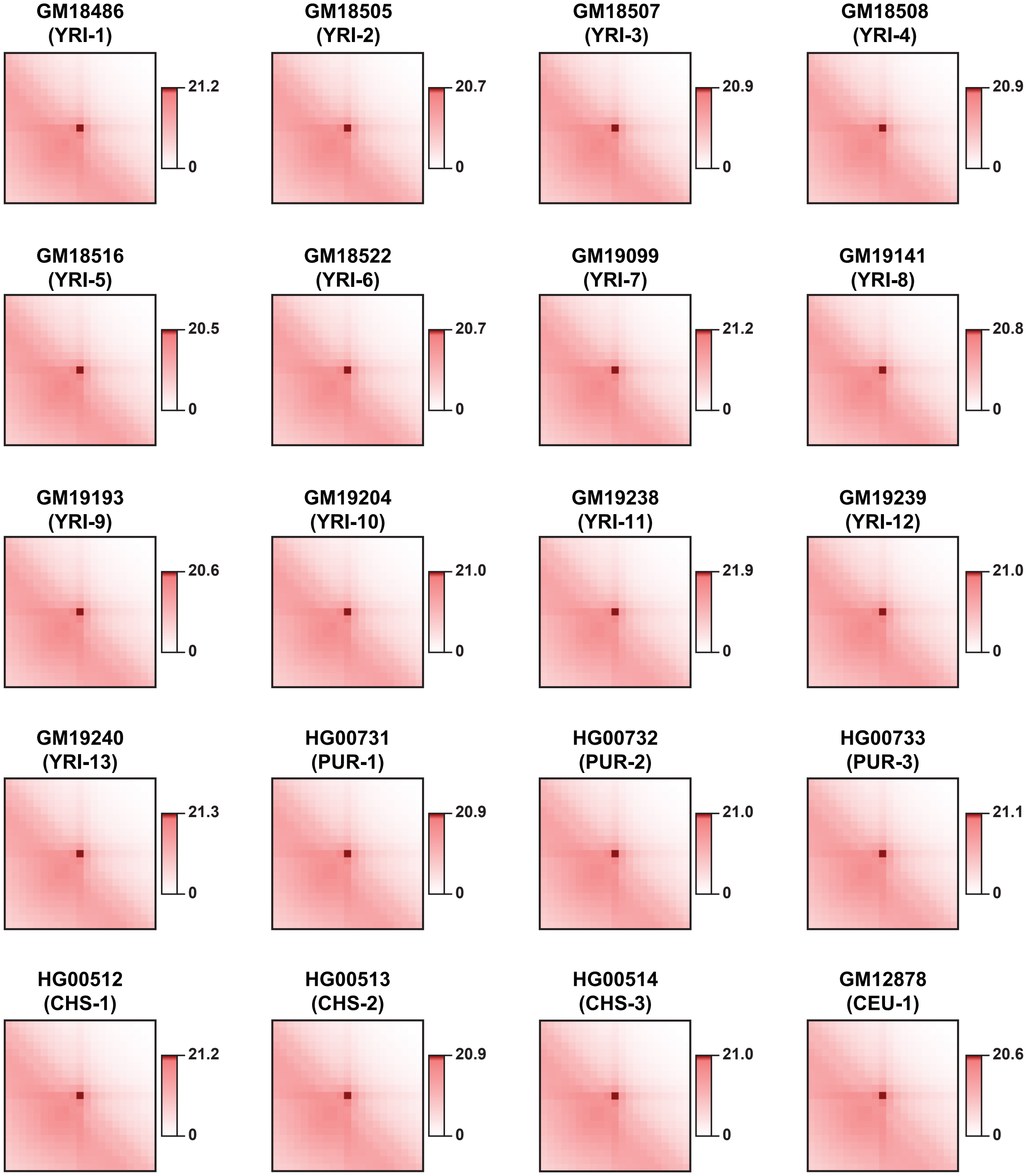


Supplemental Figure 3

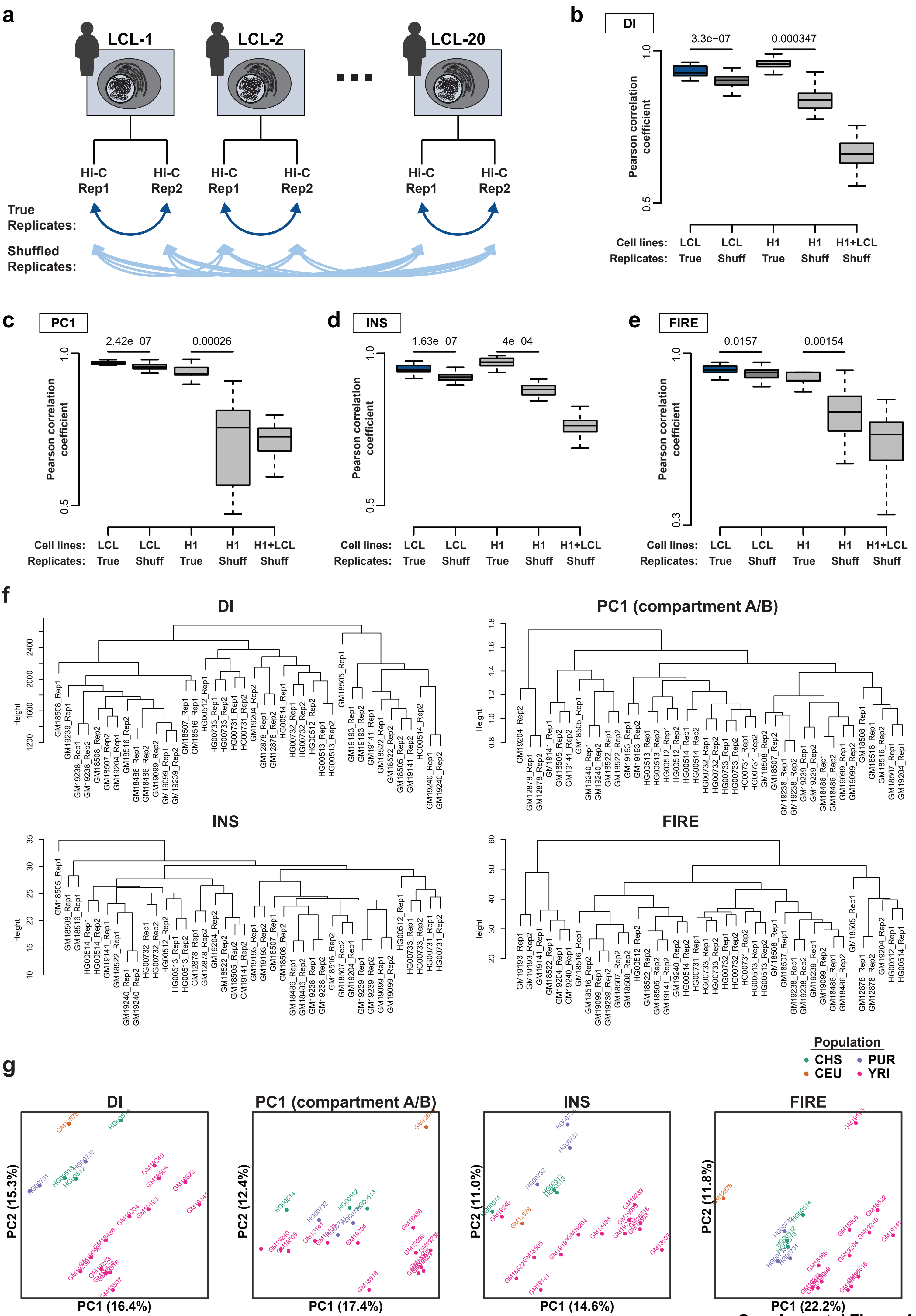


Supplemental Figure 4

a

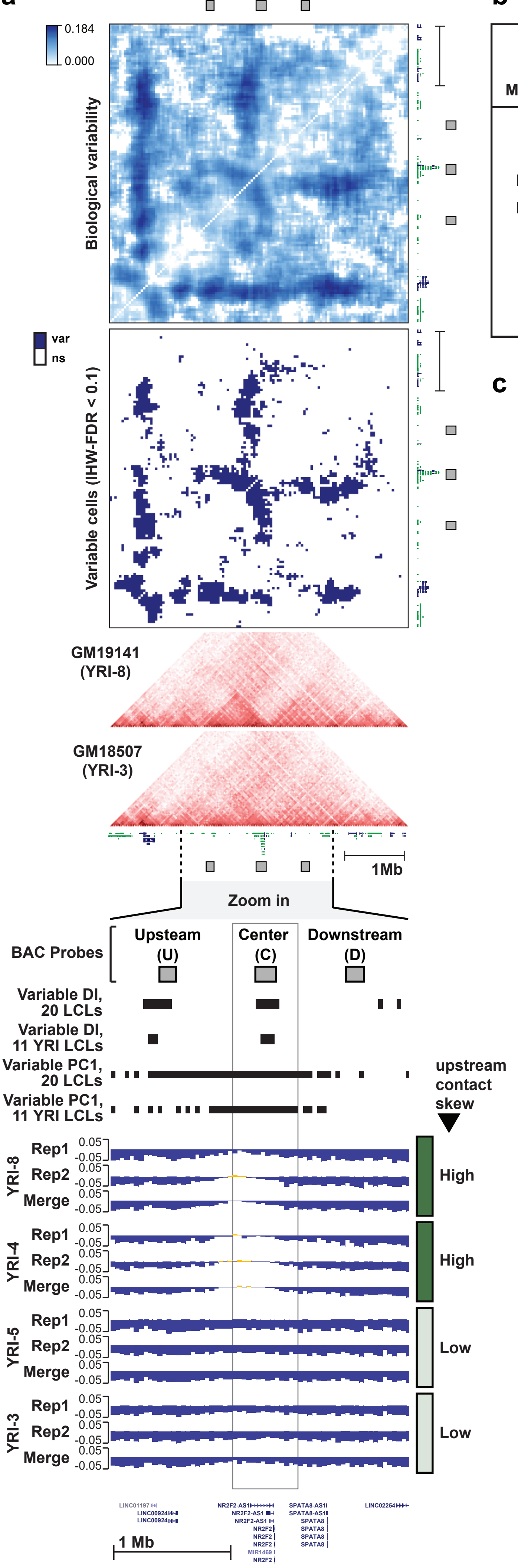


b

| Metric | discovery set | testable bins | variable bins (limma FDR<0.1) | merged regions | empirical false positives (avg 1K permutations) | empirical FDR |
|---|---|---|---|---|---|---|
| DI | 20 LCLs | 17,178 | 3,424 | 2,318 | 59.6 | 5.49% |
| DI | 11 YRI LCLs | 16,605 | 1,086 | 818 | 13.6 | 0.40% |
| FIRE | 20 LCLs | 7,276 | 3,219 | 1,993 | 29.2 | 8.92% |
| FIRE | 11 YRI LCLs | 6,321 | 327 | 255 | 24.3 | 0.75% |
| INS | 20 LCLs | 15,387 | 6,661 | 2,485 | 47.3 | 1.93% |
| INS | 11 YRI LCLs | 14,319 | 2,447 | 1,033 | 7.9 | 0.12% |
| PC1 | 20 LCLs | 52,328 | 19,128 | 7,732 | 150.6 | 2.22% |
| PC1 | 11 YRI LCLs | 52,430 | 6,777 | 3,360 | 24.2 | 0.13% |

c

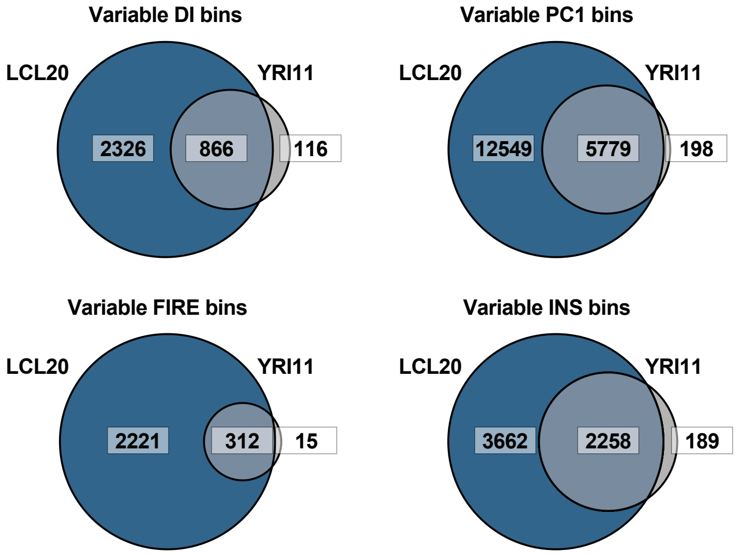


d

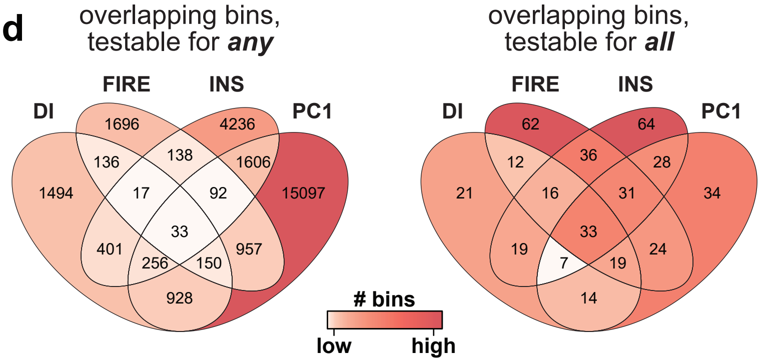


e

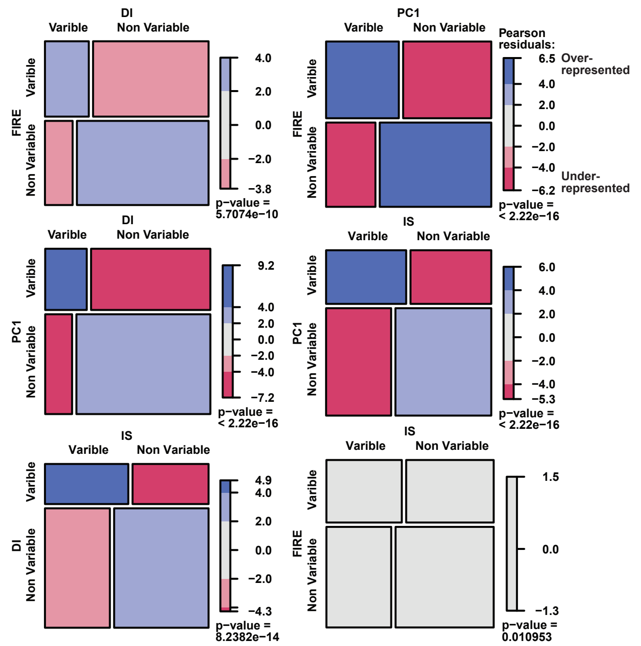


Supplemental Figure 5

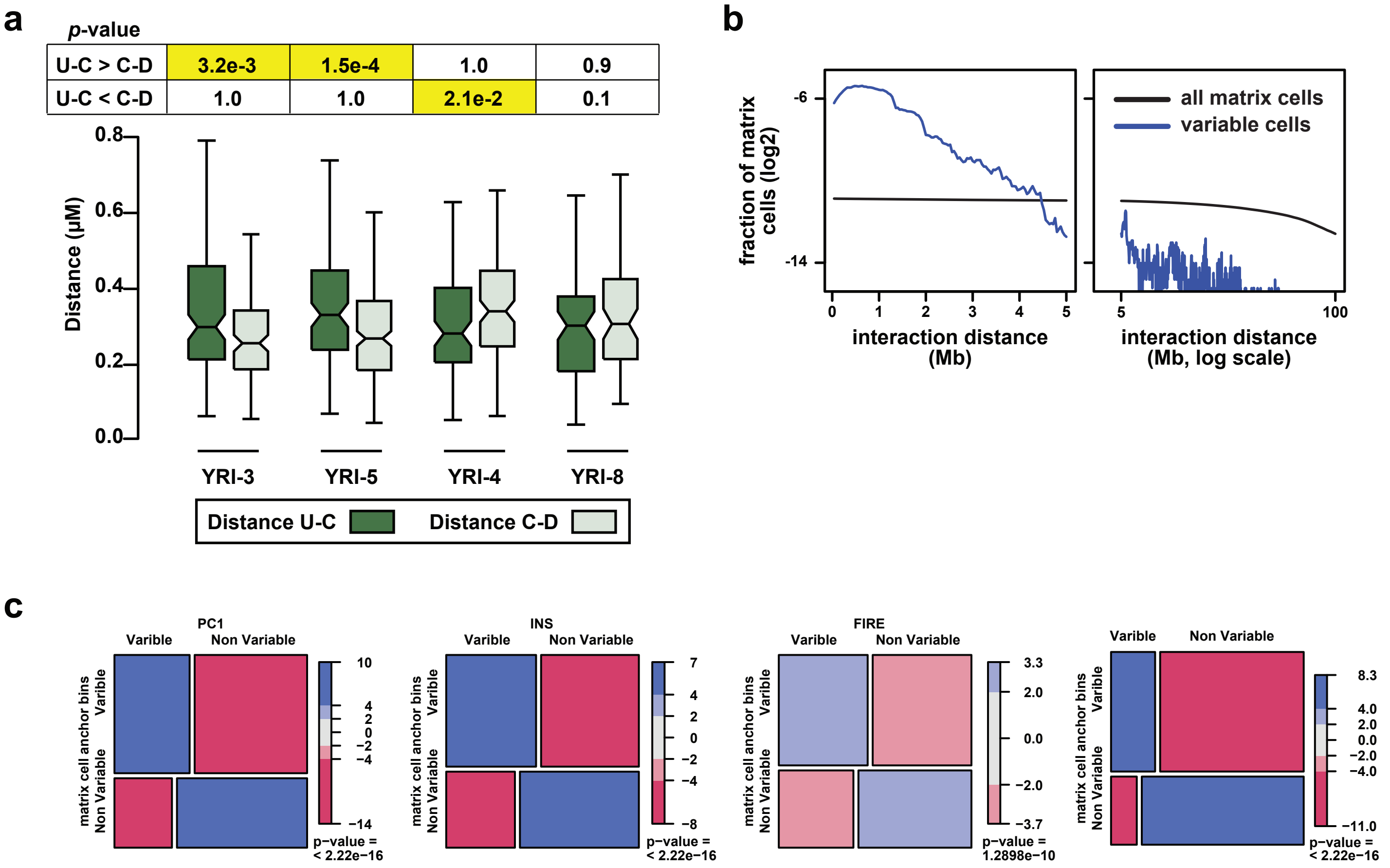


Supplemental Figure 6

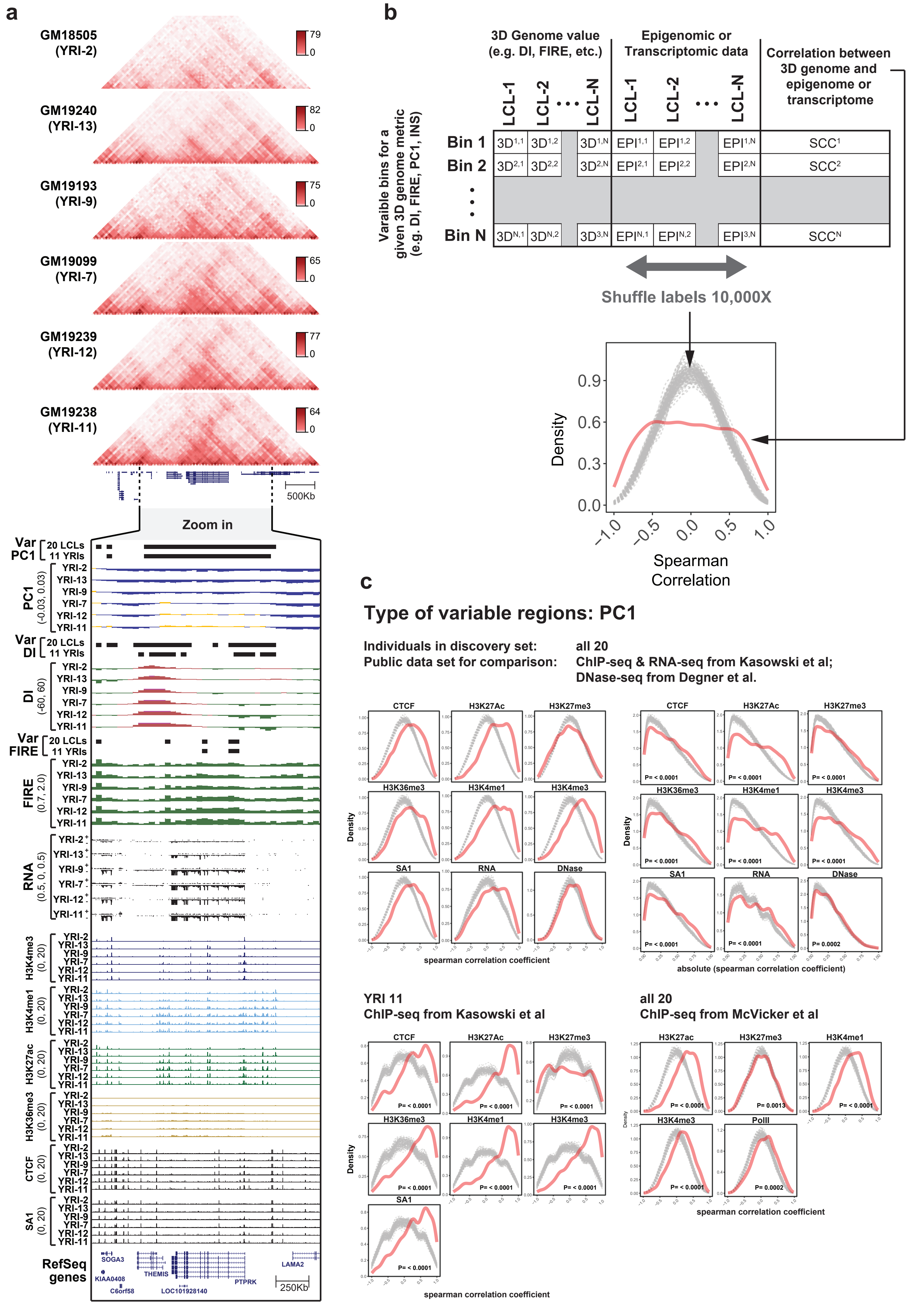


Supplemental Figure 7

a

### Type of varaible regions: DI

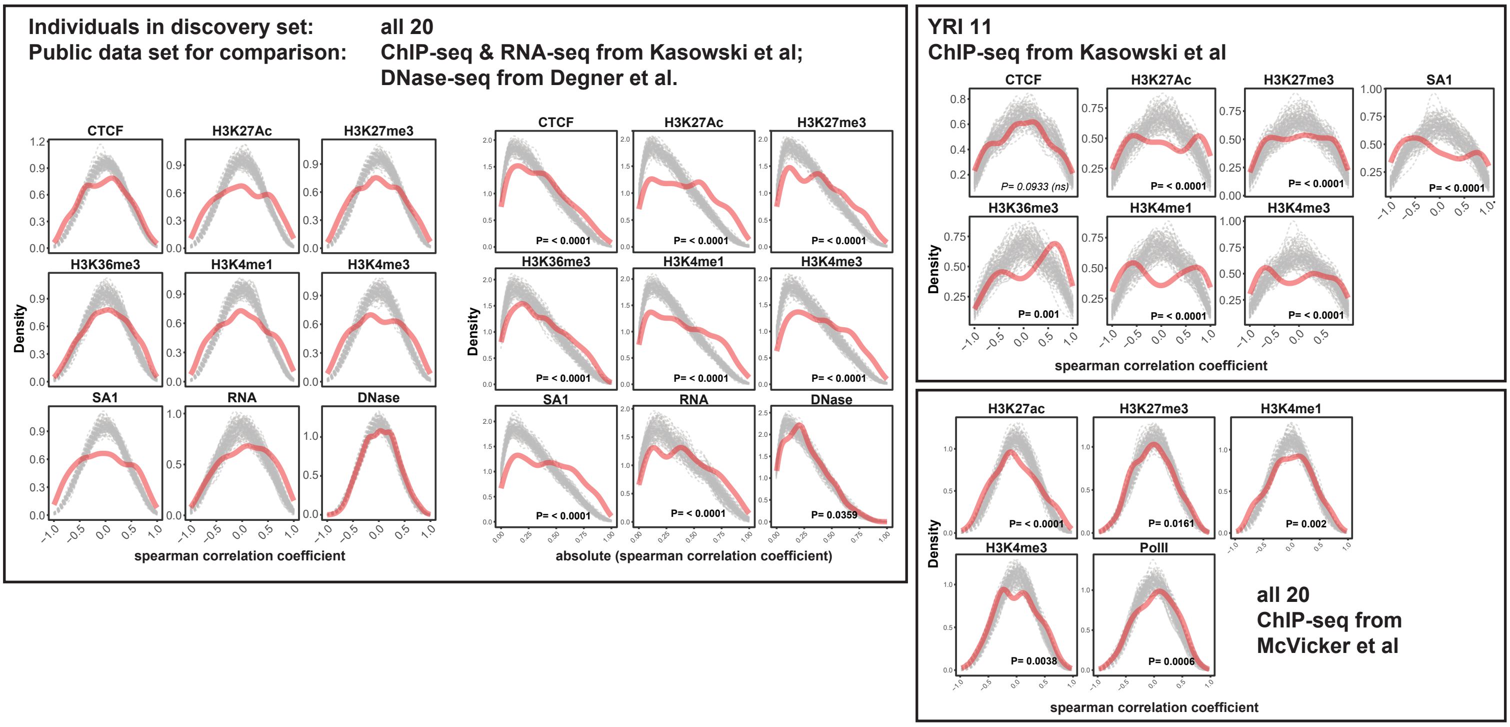


b

### Type of varaible regions: INS

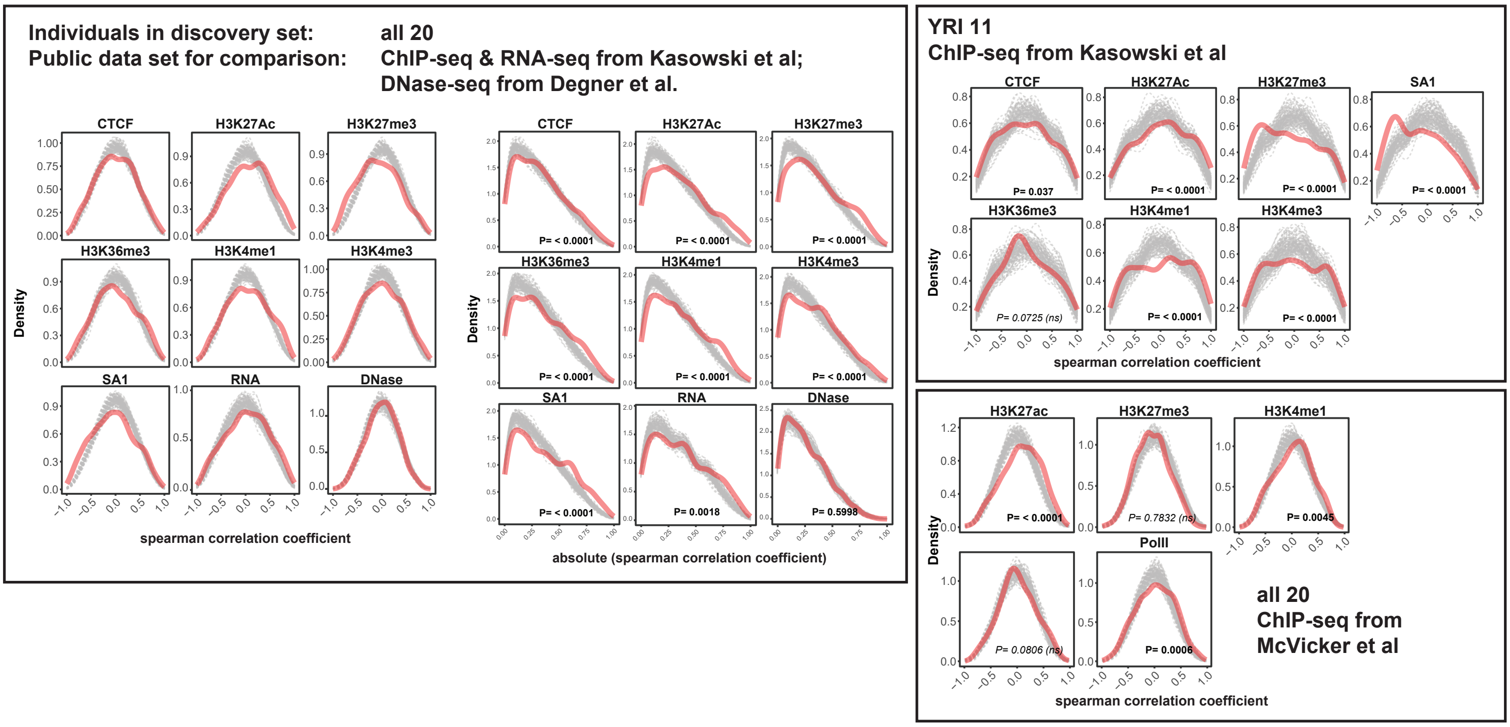

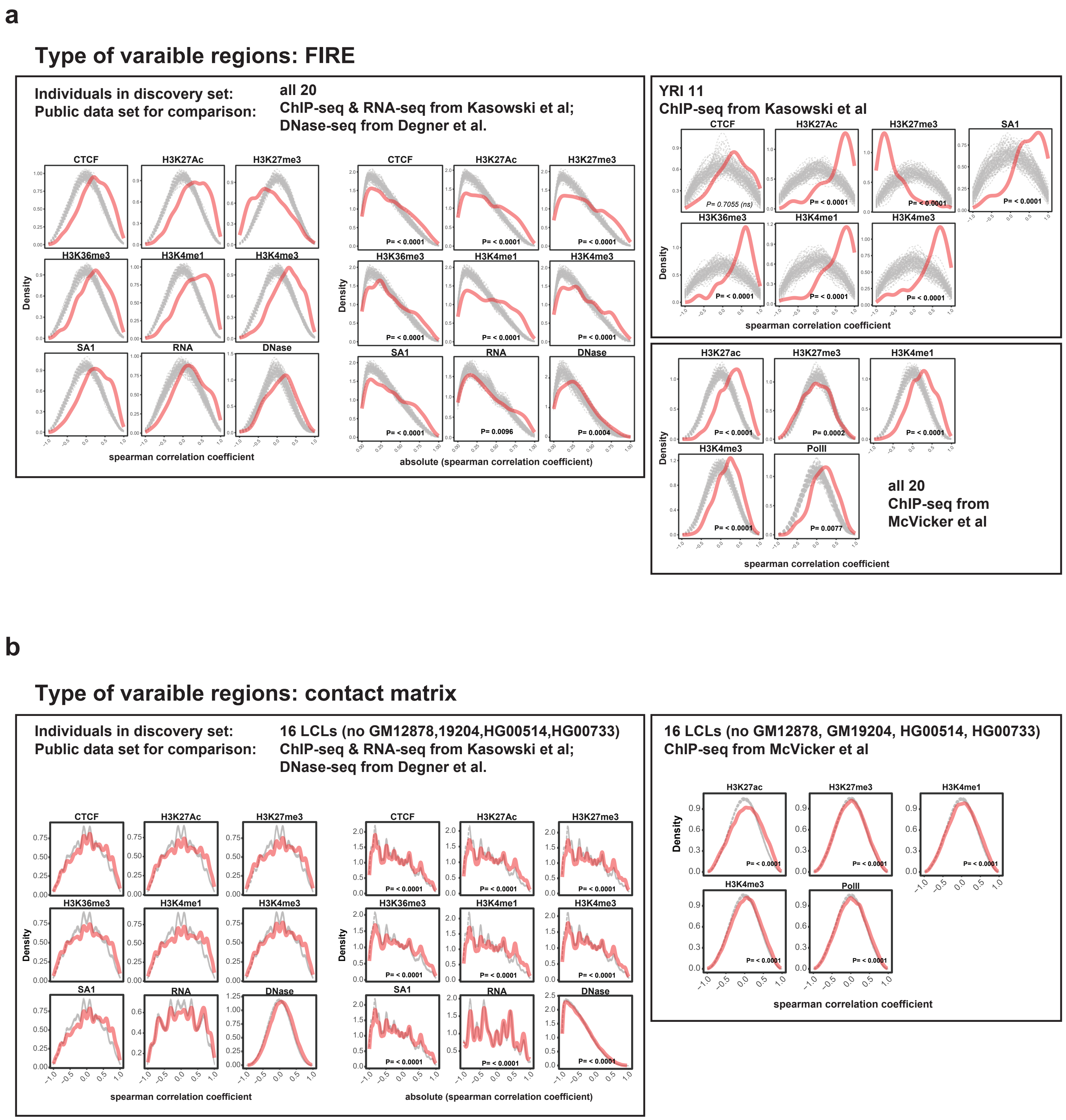


Supplemental Figure 9

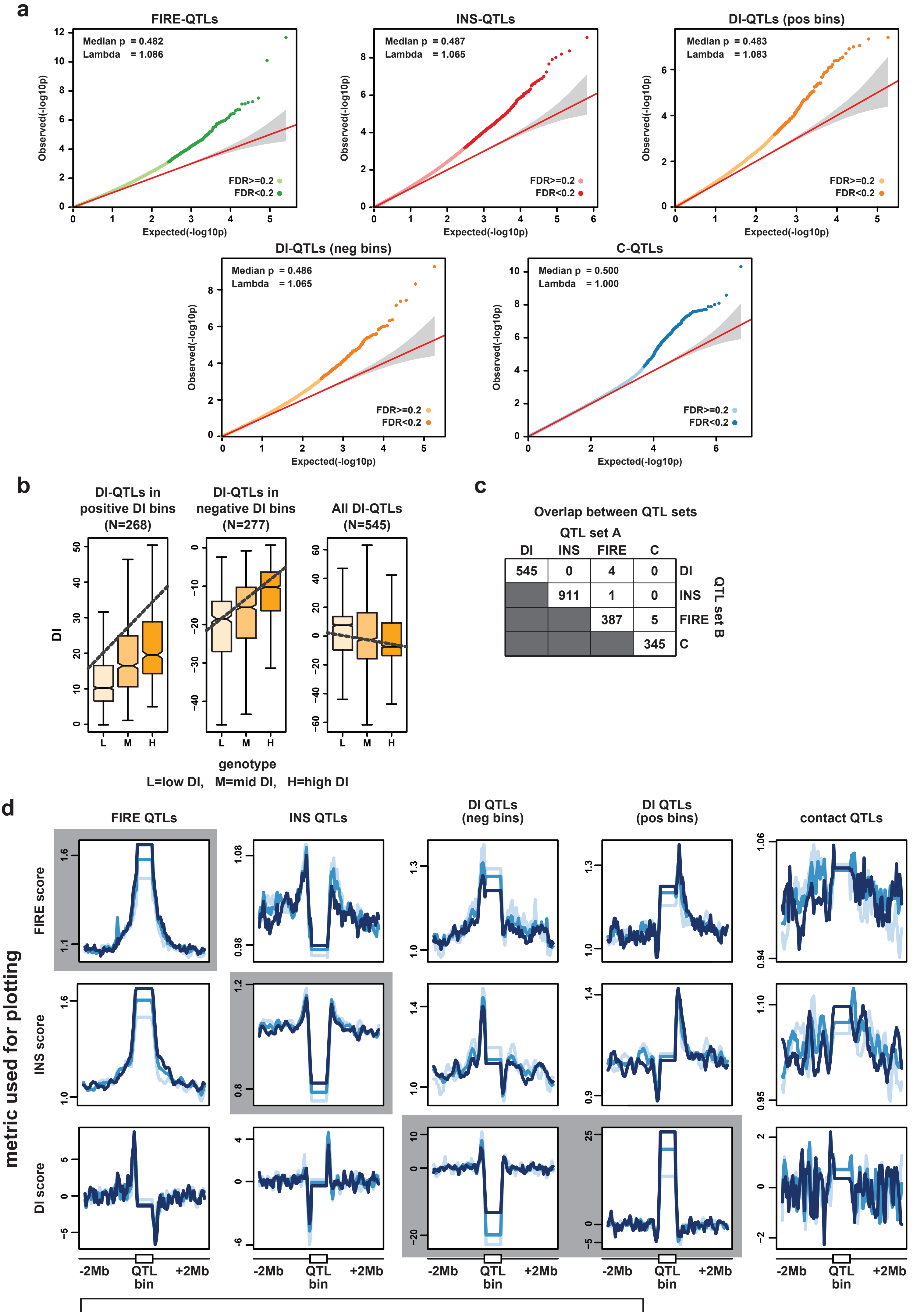


Supplemental Figure 10

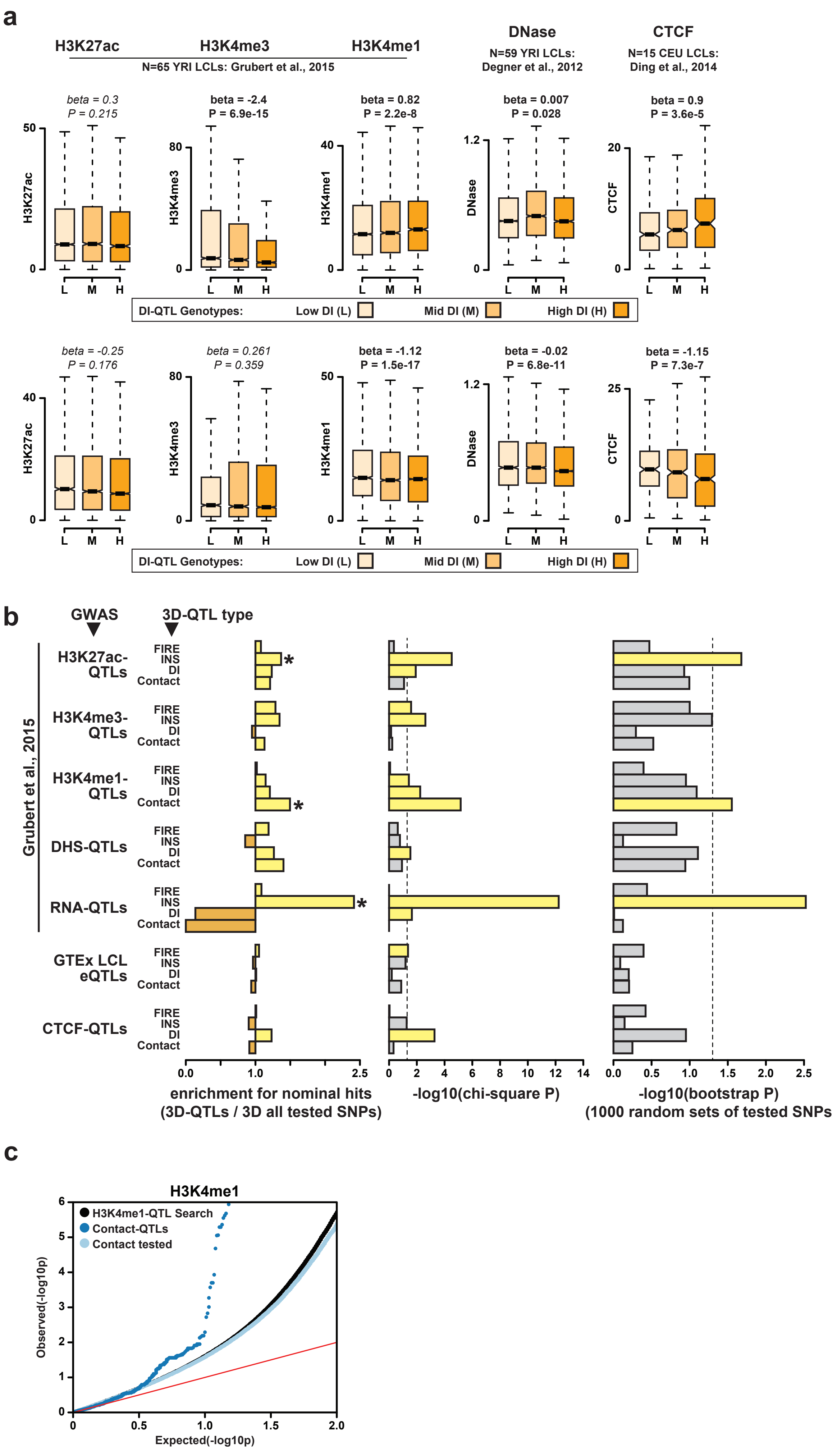


Supplemental Figure 11
